# Intra-uterine position modulates lifelong hippocampal neurogenesis and anxiety-related behavior

**DOI:** 10.64898/2026.09.29.755290

**Authors:** Fabio Grieco, Marta Vilademunt, Maryam Ali, Thibault Sprenger, Pauline van Gelderen, Hector Gallart-Ayala, Julijana Ivanisevic, Ashton Holub, June Yoo, Tracy Bedrosian, Thomas Larrieu, Nicolas Toni

## Abstract

Individual differences in vulnerability to disease are a defining feature of biology and are especially prominent in mood disorders, where susceptibility to stress varies markedly across individuals. Strikingly, such individuality also emerges in inbred mice despite genetic homogeneity and controlled environments, suggesting that early non-genomic factors can durably shape behavioral trajectories. Yet the developmental origin of this variability remains largely unknown. Here, we identify the intra-uterine position (IUP) as a potent source of inter-individual variability in emotional behavior by showing that it influences prenatal steroid exposure, hippocampal development, adult neurogenesis, and anxiety-like traits in inbred mice. At embryonic day 18.5, embryos adjacent to one or two male siblings (i.e. 1M/2M) exhibited altered brain steroid profiles, coinciding with increased proliferation of neural stem and progenitor cells in the dentate gyrus, as compared to 0M mice. Single-nucleus RNA sequencing further revealed that IUP leaves a transcriptional imprint on hippocampal development, with 1M/2M neural stem cells showing reduced stemness and activation of a neurogenic transcription program. In adulthood, 1M/2M mice presented reduced hippocampal and dentate gyrus volumes, diminished radial-glia-like cell counts, lower cell proliferation, and impaired dendritic maturation of immature neurons. Behaviorally, both juvenile (PND22) and young adult (8-week) 1M/2M mice displayed increased trait anxiety, and 1M/2M males exhibited heightened stress reactivity. *In vitro*, testosterone, but not progesterone, directly enhanced proliferation of adult rat hippocampal stem/progenitor cells, whereas corticosterone reduced it. *In vivo*, administration of the androgen receptor agonist dihydrotestosterone (DHT) to pregnant dams recapitulated IUP effects by increasing embryonic dentate gyrus (DG) proliferation in female embryos, whereas antagonism reduced progenitor proliferation in male embryos. Together, these results indicate that IUP has long-lasting effects on hippocampal development and anxiety and may underlie natural variations in trait anxiety.

## Introduction

Gaining insight into the early factors that shape individual susceptibility to mood disorders is essential for understanding why individuals exposed to apparently similar conditions can follow markedly different behavioral and neurobiological trajectories. Although the genetic background can influence an individual’s susceptibility to adverse events ultimately leading to mood disorders, twin studies indicate that non-genetic factors play an important role as well^1, 2^.

Inbred rodent studies have been instrumental to examine the causative role of non-genetic factors in the development of stress susceptibilities and have identified important contributors such as environmental enrichment, physical activity^3, 4, 5^ or maternal care^6, 7^. Such non-shared experience leads to interindividual variability in behavior and underlying biological mechanisms^8–10^. However, early life experiences are not sufficient to explain interindividual variability^11^ and may therefore involve prenatal non-shared experience. In rodents, females have 2 uterine horns in which embryos are randomly aligned and can therefore be positioned between two males, one male and one female, or two females, commonly referred to as 2M, 1M, and 0M intra-uterine positions (IUP), respectively. As hormones and other signaling molecules can be exchanged through the amniotic membranes, this spatial arrangement exposes embryos from the same litter to distinct intra-uterine hormonal milieu^12^. In particular male embryos produce testosterone earlier and at higher concentrations than females, and this hormone can diffuse to adjacent embryos^13, 14^. As a consequence, embryos located between two males are exposed to higher local androgen concentrations than embryos without adjacent male neighbors^13^. IUP therefore constitutes a naturally occurring source of non-genomic variability, capable of generating developmental differences among genetically very similar individuals sharing the same postnatal environment^15^. Consistent with this view, proximity to male embryos *in utero* has been linked to increased anogenital distance (AGD), higher body weight, increased aggression and territoriality, and reduced parental behavior in adulthood^16, 17^. Together, these observations indicate that IUP can have consequences later in life. However, little is known about the effect of IUP on brain development.

Given the role of prenatal steroid exposure in developmental programming and the central contribution of hippocampal function to emotional regulation, we hypothesized that IUP may represent an early non-genomic source of individual variability that shapes hippocampal development, and vulnerability to anxiety-related behavior.

## Results

### Intra-uterine position influences the development of brain areas associated with anxiety

IUP has been shown to influence several parameters such as body weight and AGD^16, 18, 19^. However, whether IUP also affects brain development remains unexplored. To address this question, pregnant females underwent caesarean sections on the expected day of parturition i.e., gestational day (GD) 18.5. The uterine horns were exposed, and each embryo was extracted individually and their sex and IUP were determined (Fig. 1A). Their brains were cryosectioned and stained to measure the volume of anxiety-related brain structures, including the hypothalamus, amygdala and hippocampus. In males, proximity to other males was associated with larger hypothalamus, amygdala, hippocampus and dentate gyrus whereas in females, proximity to male increased hippocampal and volumes only (Fig. 1B-I). These IUP-dependent volume differences were not associated with differences in body weight or brain weight (Suppl. Fig. 1A-D), suggesting that *in utero* male proximity influences the development of specific brain areas.

**Figure 1.**
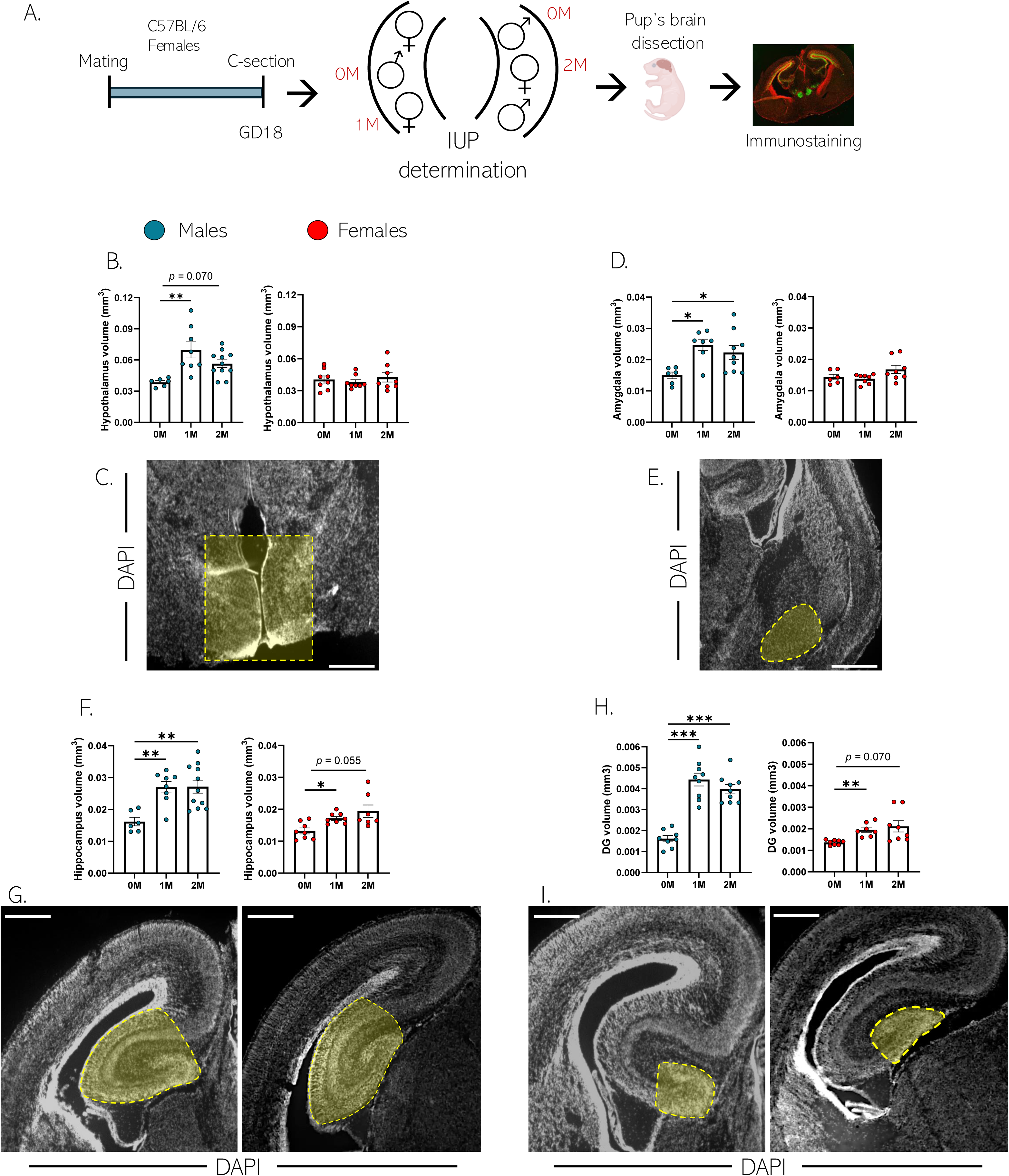
IUP influences brain development: **A.** Schematic illustration of the experimental design. **B.** Hypothalamus volume in males (left panel) and females (right panel) (**Males**: F2,22 = 4.279, p = 0.027, one-way ANOVA, n = 6, 8 and 11 mice per group; Tukey’s multiple comparisons test, 0M vs 1M: p = 0.002, 0M vs 2M: p = 0.070; 1M vs 2M: p = 0.163. **Females**: F2,21 = 0.421, p = 0.662, one-way ANOVA, n = 8 per group). **C.** Confocal image of the hypothalamus (highlighted in yellow), immunostained for DAPI (grey) (scale bar 1000µm). **D.** Amygdala volume in males (left panel) and females (right panel) (**Males**: F2,19 = 5.914, p = 0.010, one-way ANOVA, n = 6, 7 and 9 mice per group; Tukey’s multiple comparisons test, 0M vs 1M: p = 0.011, 0M vs 2M: p = 0.033; 1M vs 2M: p = 0.371. **Females**: F2,19 = 2.803, p = 0.086, one-way ANOVA, n = 6, 8 and 8 per group). **E.** Confocal image of the amygdala (highlighted in yellow), immunostained for DAPI (grey) (scale bar 1000µm). **F.** Hippocampus volume in males (left panel) and females (right panel) (**Males**: F2,22 = 8.677, p = 0.002, one-way ANOVA, n = 6, 8 and 11 mice per group; Tukey’s multiple comparisons test, 0M vs 1M: p = 0.005, 0M vs 2M: p = 0.002; 1M vs 2M: p = 0.997. **Females**: F2,10 = 5.865, p = 0.020, Brown-Forsythe ANOVA test, n = 8 per group; Dunnett’s T3 multiple comparisons test, 0M vs 1M: p = 0.011, 0M vs 2M: p = 0.055; 1M vs 2M: p = 0.644). **G.** Confocal image of the dorsal (left panel) and ventral (right panel) hippocampus (highlighted in yellow), immunostained for DAPI (grey) (scale bar 1000µm). **H.** Dentate gyrus volume in males (left panel) and females (right panel) (**Males**: F2,23 = 37.80, p = < 0.001, one-way ANOVA, n = 8 and 9 mice per group; Tukey’s multiple comparisons test, 0M vs 1M: p = < 0.001, 0M vs 2M: p = < 0.001; 1M vs 2M: p = 0.372. **Females**: F2,9.85 = 8.574, p = 0.002, Brown-Forsythe ANOVA test, n = 8, 7 and 8 per group; Dunnett’s T3 multiple comparisons test, 0M vs 1M: p = 0.006, 0M vs 2M: p = 0.070; 1M vs 2M: p = 0.928). **I.** Confocal image of the dorsal (left panel) and ventral (right) hippocampus (highlighted in yellow), immunostained for DAPI (grey) (scale bar 1000µm). Histograms show average ± SEM, * p<0.05, ** p<0.01, ***p<0.001, ns = not significant.

Since the dentate gyrus displayed the most important volume change with IUP, we further investigated neurogenesis in this area, using molecular markers for neural stem cells (HOPX) neural progenitors (Tbr2), immature neurons (NeuroD1) and cell proliferation (Ki-67, Fig. 2A). In both males and females, the proximity to males *in utero* was linked to fewer HOPX^+^ and Tbr2^+^ cells. Although IUP did not affect overall cellular proliferation, the presence of a single adjacent male was sufficient to increase proliferation within both cell types (Fig. 2B-H). Analysis of immature neurons further showed that 2M embryos displayed a higher number of NeuroD1^+^ cells compared with 0M individuals, regardless of sex (Fig. 2I). Taken together, these findings indicate that IUP is associated with selective changes in late-embryonic brain development and with an altered neurogenic profile within the dentate gyrus of the hippocampus.

**Figure 2.**
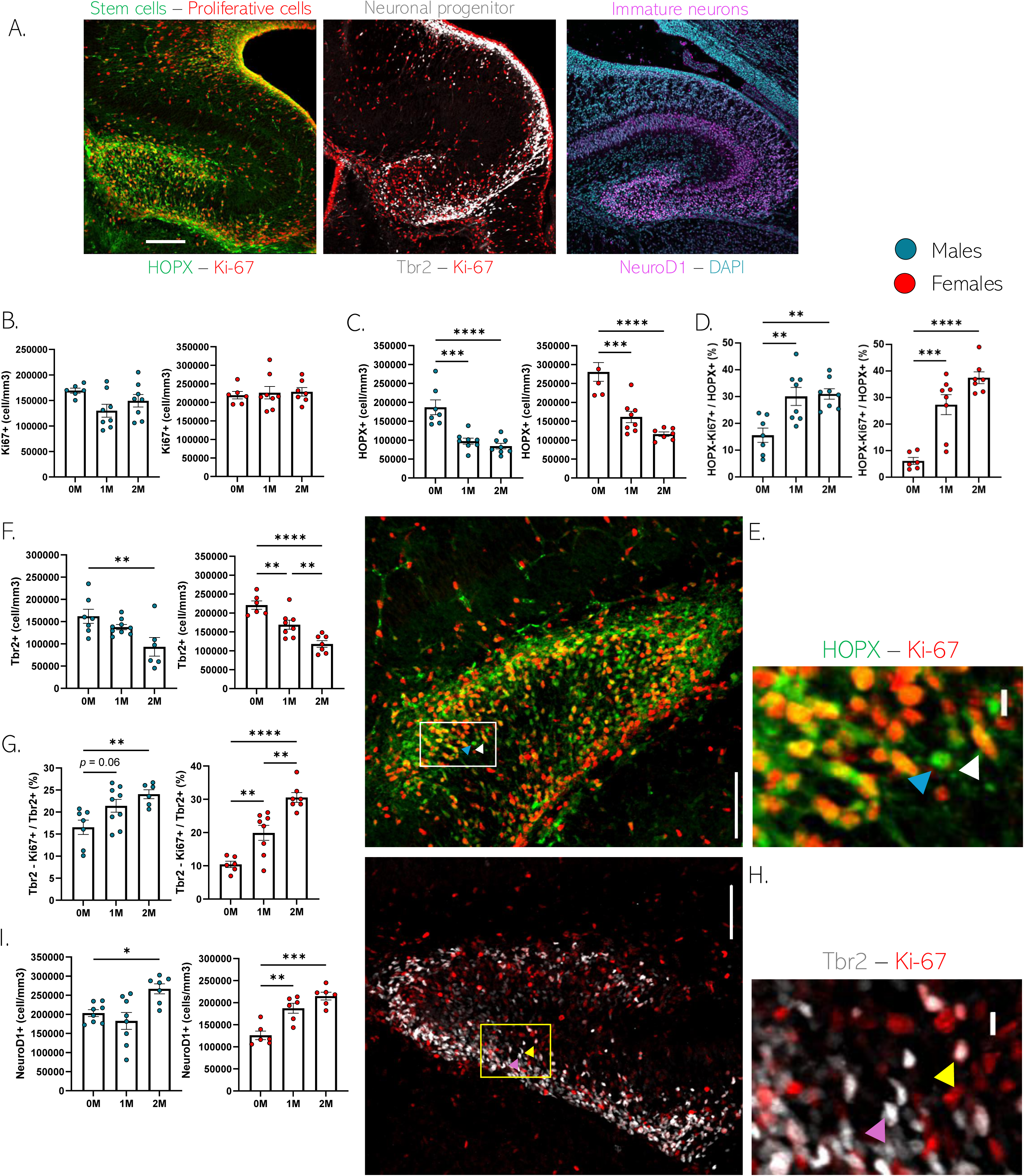
IUP influences embryonic neurogenesis in the DG: **A.** Confocal maximal projection of DG slices immunostained for stem cells, proliferative cells (HOPX in green and Ki-67 in red; left panel), neural progenitors (Tbr2 in grey, central panel) and immature neurons (NeuroD1 in magenta; right panel) in embryos at GD 18.5 (scale bar 100µm). **B.** Quantification of Ki-67^+^ cells in males (left panel) and females (right panel) (**Males**: F2,19 = 2.853, p = 0.082, one-way ANOVA, n = 7, 8 and 8 mice per group. **Females**: F2,18 = 0.108, p = 0.897, one-way ANOVA, n = 6, 8 and 7 per group). **C.** Quantification of HOPX^+^ cells in males (left panel) and females (right panel) (**Males**: F2,20 = 20.30, p = <0.0001, one-way ANOVA, n = 7, 8 and 8 mice per group; Tukey’s multiple comparisons test, 0M vs 1M: p = 0.0001, 0M vs 2M: p = <0.0001; 1M vs 2M: p = 0.7154. **Females**: F2,18 = 16.26, p = <0.0001, one-way ANOVA, n = 6, 8 and 7 per group; Tukey’s multiple comparisons test, 0M vs 1M: p = 0.0002, 0M vs 2M: p = <0.0001; 1M vs 2M: p = 0.1259). **D.** Percentage of HOPX^+^-Ki-67^+^ cells over the total amount of HOPX^+^ cells (**Males**: F2,20 = 9.451, p = 0.0013, one-way ANOVA, n = 7, 8 and 8 mice per group; Tukey’s multiple comparisons test, 0M vs 1M: p = 0.004, 0M vs 2M: p = 0.0023; 1M vs 2M: p = 0.9968. **Females**: F2,18 = 27.66, p = <0.0001, one-way ANOVA, n = 6, 8 and 7 per group; Tukey’s multiple comparisons test, 0M vs 1M: p = 0.0002, 0M vs 2M: p = <0.0001; 1M vs 2M: p = 0.0501). **E.** Representative confocal microscopy image of the dentate gyrus immunostained for HOPX and HOPX-Ki-67-expressing cells (HOPX^+^ highlighted with blue arrow heads and HOPX^+^-Ki-67^+^ highlighted with white arrow heads; scale bar 100µm and 10µm in the inset). **F.** Quantification of Tbr2^+^ cells in males (left panel) and females (right panel) (**Males**: F2,19 = 5.631, p = 0.012, one-way ANOVA, n = 7, 9 and 6 mice per group; Tukey’s multiple comparisons test, 0M vs 1M: p = 0.4111, 0M vs 2M: p = 0.0095; 1M vs 2M: p = 0.0858. **Females**: F2,18 = 20.82, p = <0.0001, one-way ANOVA, n = 6, 8 and 7 per group; Tukey’s multiple comparisons test, 0M vs 1M: p = 0.0095, 0M vs 2M: p = <0.0001; 1M vs 2M: p = 0.0079). **G.** Percentage of Tbr2^+^-Ki-67^+^ cells over the total amount of Tbr2^+^ cells (**Males**: F2,19 = 6.195, p = 0.0085, one-way ANOVA, n = 7, 9 and 6 mice per group; Tukey’s multiple comparisons test, 0M vs 1M: p = 0.0599, 0M vs 2M: p = 0.0077; 1M vs 2M: p = 0.4279. **Females**: F2,18 = 29.38, p = <0.0001, one-way ANOVA, n = 6, 8 and 7 per group; Tukey’s multiple comparisons test, 0M vs 1M: p = 0.044, 0M vs 2M: p = <0.0001; 1M vs 2M: p = 0.011). **H.** Representative confocal microscopy image of the dentate gyrus immunostained for Tbr2 and Tbr2^+^-Ki-67^+^ cells (Tbr2^+^ highlighted with pink arrows and Tbr2^+^-Ki-67^+^ highlighted with yellow arrows; scale bar 100µm and 10µm in the inset). **I.** Quantification of NeuroD1 positive cells in males (left panel) and females (right panel) (**Males**: F2,20 = 7.116, p = 0.005, one-way ANOVA, n = 9, 8 and 7 mice per group; Tukey’s multiple comparisons test, 0M vs 1M: p = 0.633, 0M vs 2M: p = 0.031; 1M vs 2M: p = 0.078. **Females**: F2,15 = 20.62, p = <0.001, one-way ANOVA, n = 6 per group; Tukey’s multiple comparisons test, 0M vs 1M: p = 0.002, 0M vs 2M: p = < 0.001; 1M vs 2M: p = 0.159). Histograms show average ± SEM, * p<0.05, ** p<0.01, ***p<0.001, ****p<0.0001 ns = not significant.

### Intra-uterine position is associated with a pro-neurogenic transcriptional program in embryonic hippocampal neural stem cells

To gain insight into the transcriptional changes associated with IUP-dependent alterations in hippocampal neurogenesis, we performed single-nucleus gene expression analysis of hippocampal tissue from embryos at GD18.5. We profiled over 44,320 nuclei after QC filtering (Suppl. Fig. 2A) which represented all major hippocampal cell types, as well as nearby cortical neuronal subtypes (Fig. 3A). Notably, we observed populations of neural stem cells identified by *Nes*, *Pax6*, and *Hopx* expression, intermediate progenitors defined by *Eomes* (TBR2), and immature DG neurons identified by *Neurod1* and *Prox1* expression. We noted that neural stem cells contained four subgroups: cycling NSCs expressing *Mki67* and *Top2a* and non-cycling NSCs lacking those markers, as well as small lineage committed subgroups expressing neurogenic and gliogenic markers (Fig. 3B).

**Figure 3:**
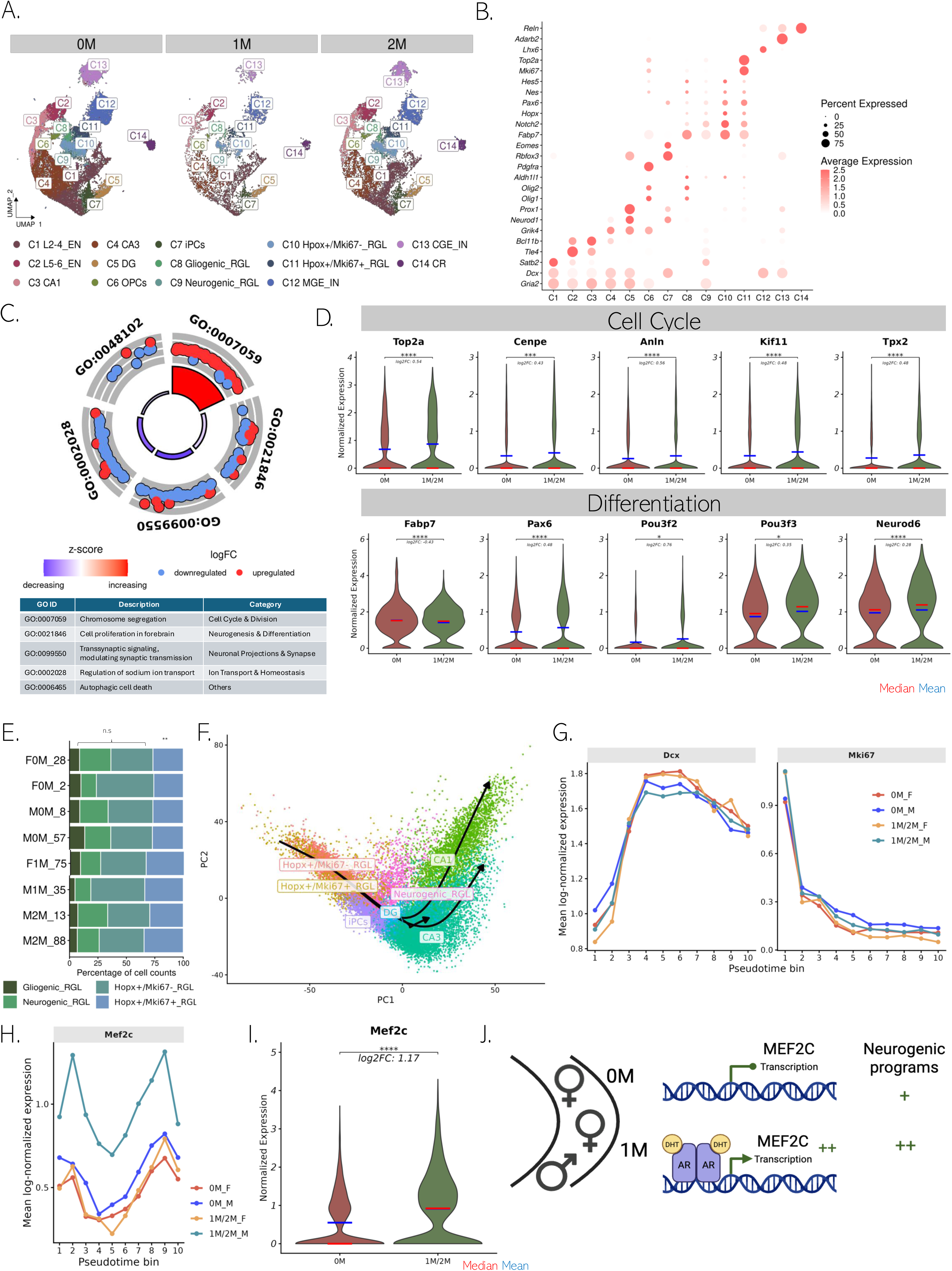
IUP drives gene expression differences in neural stem cells: **A.** UMAP plots of single-nucleus RNA-sequencing data identifying cell clusters present in 0M, 1M, and 2M mice. **B.** Expression of cell type marker genes across clusters. **C.** Representative GO terms identified from differential gene expression analysis comparing 1M/2M vs. 0M cells within NSCs. **D.** Expression of representative genes differentially expressed among IUP groups with sex included as a covariate (MAST, Padj < 0.05). **E.** Proportion of NSC subtypes by mouse (quasibinomial GLM, BH-adjusted p-value = 0.0068 for Hopx^+^/Mki67^+^ RGL cluster). **F.** Slingshot pseudotime trajectories demonstrating the progression from cycling NSC to mature hippocampal neuronal subtypes. **G.** *Mki67* and *Dcx* show expected expression dynamics from early to late pseudotime, supporting a cycling-to-neurogenic differentiation trajectory. Statistical significance was assessed using tradeSeq startVsEndTest. (Wald test, adjusted p-value < 2.2 × 10^-16^). **H.** *Mef2c* is consistently upregulated in 1M/2M vs. 0M along pseudotime (tradeSeq condition Test, BH-adjusted p-value = 3.55 × 10^-4^). **I.** Within NSCs, *Mef2c* is significantly upregulated, particularly in 1M/2M males (MAST, padj <0.05). **J.** Theoretical model for the effect of IUP on *Mef2c* expression and subsequent neurogenesis through AR responsiveness.

To investigate potential gene expression changes driven by IUP, we performed differential gene expression analysis within cell types. Specifically, we compared embryos with (1M/2M) vs. without (0M) proximity to a male embryo, including sex as a covariate. NSCs had the highest number of differentially expressed genes (DEGs) compared to other hippocampal cell types (Suppl. Fig. 2B). Within NSCs, we identified 374 total DEGs enriched in gene ontology pathways related to the cell cycle, mitosis, and cell growth and differentiation (Fig. 3C; Suppl. Fig. 2C). Consistent with our histological data, we found that the presence of an adjacent male was associated with increased expression of proliferation-related genes, including *Top2a* and *Cenpe*, reduced expression of stemness-associated markers such as *Fabp7*, and activation of a neurogenic transcriptional program, including *Pax6, Pou3f2/3 and Neurod6*, suggesting a shift in the neurogenic trajectory (Fig. 3D). Additional IUP-regulated genes, including Apoe, Gm2a, Syp and Ptgds, further supported the presence of a distinct transcriptional state in 1M/2M NSCs compared with 0M NSCs (Suppl. Fig. 4B). The proportion of cells belonging to each NSC subtype was mostly unchanged across IUP, but there was an increase in the proportion of Hopx^+^-Mki67^+^ NSCs in 1M/2M, consistent with our histological data (Fig. 3E, Fig.2).

As IUP and sex are closely related concepts, we performed a second differential gene expression analysis directly comparing the effects of IUP (1M/2M vs. 0M) or sex (M vs. F) within NSCs. In total, 87 DEGs (12.7%) overlapped between the two comparisons (Suppl. Fig. 3A) and 90.8% of overlapping genes were concordant in their directionality (Suppl. Fig. 3B). Common DEGs were enriched in GO terms related to development and axonogenesis, whereas genes specific to the sex comparison were related to ion and signal transduction pathways. Interestingly, genes related to cell division and mitosis were specific for the IUP comparison (Suppl. Fig. 3C). Representative sex-specific DEGs included Hopx, Notch1, Sox2 and Foxo1 (Suppl. Fig. 3D), whereas representative common DEGs included Neurod6, Pou3f2, Fabp7 and Mef2c (Suppl. Fig. 3E). Together, this analysis suggests that although IUP and sex share a partially overlapping transcriptional signature, the IUP-associated program includes a distinct component related to cell division and mitotic activity within embryonic NSCs. Given the coordinated transcriptional shift toward cell division and a pro-neurogenic state in 1M/2M IUP, we performed pseudotime trajectory analysis to model gene regulation as cells differentiate from NSCs to mature hippocampal neuronal subtypes. We identified three lineages underlying the differentiation from NSCs to CA1 pyramidal neurons, CA3 pyramidal neurons, and DG granule neurons (Fig. 3F). Regardless of IUP, we observed a similar distribution of cells across pseudotime, suggesting both groups are progressing through the same differentiation program, with expected genes like *Mki67* and *Dcx* varying across pseudotime in both groups (Fig. 3G). However, IUP was associated with changes in the level of gene expression over pseudotime. Among genes displaying IUP-dependent expression dynamics over pseudotime, *Mef2c* emerged as the most consistently altered between 0M and 1M/2M cells across differentiation stages (Fig. 3H).

This was notable because *Mef2c* is a crucial transcription factor involved in neurodevelopment, which promotes NSC differentiation into neurons and enhances survival, maturation and synaptic development. It exhibits dose-dependent roles in hippocampal neurogenesis where it influences neuronal commitment^20, 21^. Interestingly, *Mef2c* is tied to androgen-dependent transcriptional regulation, with androgen receptor binding sites located within the gene itself. Prenatal androgens such as DHT can upregulate *MEF2C* in human neural stem cells, influencing neurogenesis and cell differentiation^22^. Although we did not detect IUP-dependent changes in androgen receptor or other steroid hormone receptor expression in our dataset (Suppl. Fig. 4A) *Mef2c* expression was strongly upregulated in 1M/2M NSCs compared with 0M NSCs, particularly in males (Fig. 3I). These findings identify *Mef2c* as a candidate transcriptional mediator linking IUP-associated hormonal differences to altered neurogenic trajectories during embryonic development (Fig. 3J). Because LINE-1 retrotransposons have been implicated in neural stem cell differentiation, neuronal lineage commitment, and steroid hormone responsiveness^23–26^, and have been proposed as a genomic mechanism linking environmental exposures to inter-individual differences in brain function and behavior^27^, we also assessed LINE-1 expression in our dataset. Using SoloTE, to quantify transposable element expression from single-nucleus RNA-sequencing data, we observed increased expression of L1MdA and L1MdT, evolutionarily younger LINE-1 families still capable of retrotransposition, in the neurogenic NSC subcluster compared with cycling or gliogenic NSCs (Suppl. Fig. 4C-D). However, we detected no effect of IUP on LINE-1 expression and no IUP-dependent differences in the expression of known LINE-1 regulators in NSCs, including Dnmt1, Dnmt3b, Trim28, Setdb1, Yy1, and Sox2 (Suppl. Fig. 4E). These results suggest that, although LINE-1 expression is associated with neurogenic NSC states, variation in uterine position is not sufficient to alter LINE-1 expression dynamics at the transcriptional level. Future studies could address whether IUP influences LINE-1 insertions at the genomic level. Together, these transcriptional analyses confirm our histological observations that IUP modifies gene expression in NSC and in particular proliferation-related genes. They further identify *Mef2c* as the most consistently IUP-altered gene, potentially linking altered neurogenesis to hormonal differences.

### Intra-uterine position shapes anxiety-like behavior and stress reactivity

Exposure to prenatal stress or hormonal fluctuations can contribute to the development of neuropsychiatric disorders^28–30^. To test whether IUP-related effects persist postnatally, influencing anxiety-related behaviors, we designed a longitudinal study. Embryos of 18.5 gestational days were delivered by cesarean, identified for sex and IUP and fostered to CD1 dams according to their IUP classification. They were then evaluated for anxiety at 3-and 8-weeks of age, immediately followed by resilience to an acute stress. They were then perfused at 10 weeks of age for histological assessment (Fig. 4A).

**Figure 4.**
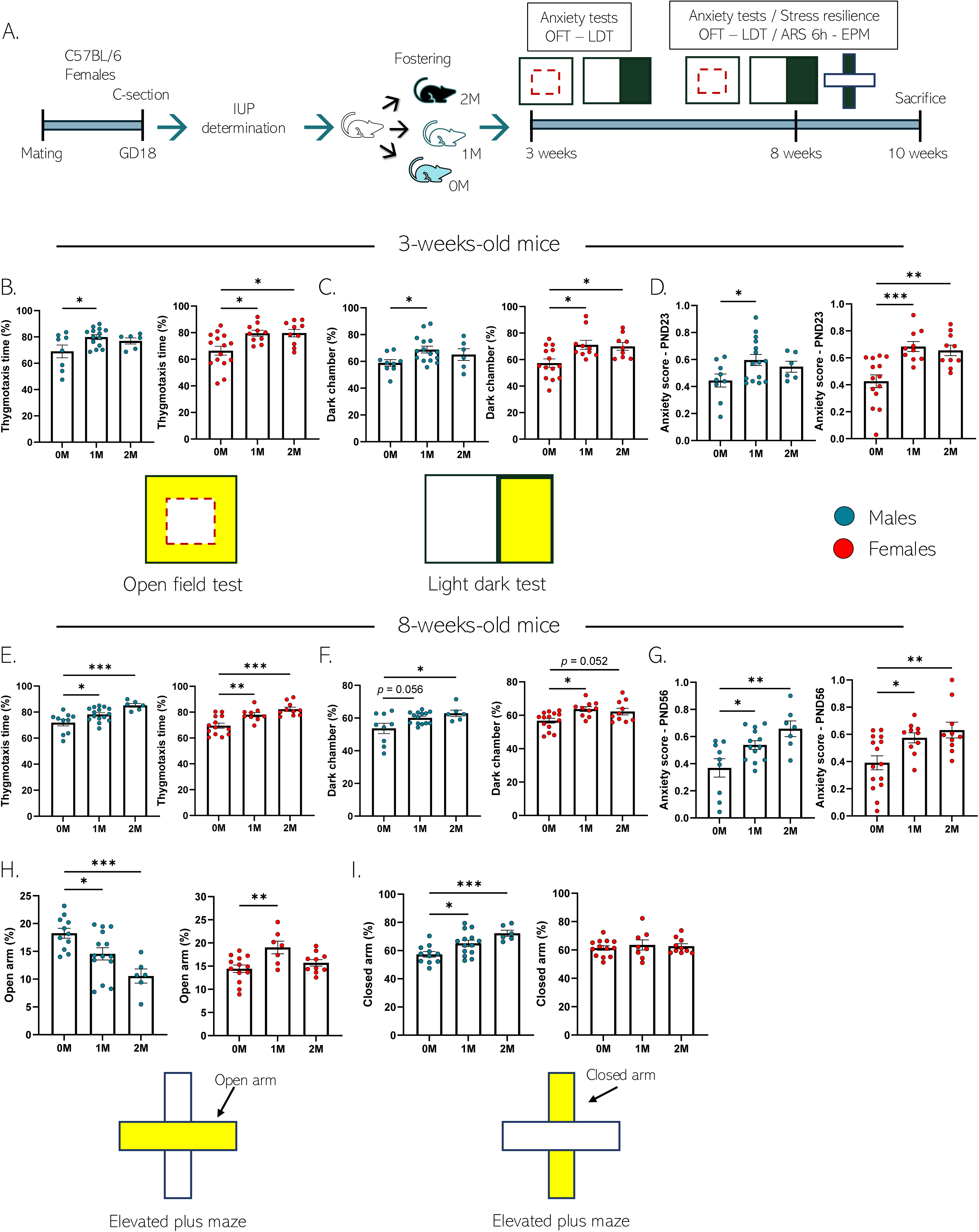
IUP influences trait anxiety in young and adult mice: **A.** Schematic illustration of the experimental design for anxiety-related behavioral assessment. **B.** Percentage of time spent in thigmotaxis in an OFT at PND22 by males (left panel) and females (right panel) (**Males**: F2,25 = 4.320, p = 0.024, one-way ANOVA, n = 8, 14 and 6 mice per group; Tukey’s multiple comparisons test, 0M vs 1M: p = 0.019, 0M vs 2M: p = 0.189; 1M vs 2M: p = 0.790. **Females**: F2,32 = 6.699, p = 0.004, one-way ANOVA, n = 15, 10 and 10 per group; Tukey’s multiple comparisons test, 0M vs 1M: p = 0.013, 0M vs 2M: p = 0.011; 1M vs 2M: p = >0.999). **C.** Percentage of time spent in the dark chamber in a LDT at PND23 by males (left panel) and females (right panel) (**Males**: F2,27 = 3.249, p = 0.049, one-way ANOVA, n = 9, 15 and 6 mice per group; Tukey’s multiple comparisons test, 0M vs 1M: p = 0.043, 0M vs 2M: p = 0.421; 1M vs 2M: p = 0.686. **Females**: F2,30 = 6.095, p = 0.006, one-way ANOVA, n = 14, 10 and 9 per group; Tukey’s multiple comparisons test, 0M vs 1M: p = 0.012, 0M vs 2M: p = 0.027; 1M vs 2M: p = 0.971). **D.** Anxiety score derived from normalized data from LDT and OFT in males (left panel) and females (right panel) (**Males**: F2,27 = 6.104, p = 0.048, one-way ANOVA, n = 9, 15 and 6 mice per group; Tukey’s multiple comparisons test, 0M vs 1M: p = 0.049, 0M vs 2M: p = 0.386; 1M vs 2M: p = 0.759. **Females**: F2,31 = 11.321, p = <0.001, one-way ANOVA, n = 14, 10 and 10 per group; Tukey’s multiple comparisons test, 0M vs 1M: p = <0.001, 0M vs 2M: p = 0.002; 1M vs 2M: p = 0.902). **E.** Percentage of time spent in thigmotaxis in an OFT at PND55 by males (left panel) and females (right panel) (**Males**: F2,28 = 9.574, p = <0.001, one-way ANOVA, n = 10, 15 and 6 mice per group; Tukey’s multiple comparisons test, 0M vs 1M: p = 0.036, 0M vs 2M: p = <0.001; 1M vs 2M: p = 0.057. **Females**: F2,29 = 13.28, p = <0.001, one-way ANOVA, n = 13, 9 and 10 per group; Tukey’s multiple comparisons test, 0M vs 1M: p = 0.008, 0M vs 2M: p = <0.001; 1M vs 2M: p = 0.305). **F.** Percentage of time spent in the dark chamber in a LDT at PND56 by males (left panel) and females (right panel) (**Males**: F2,13.79 = 4.157, p = 0.039, Brown-Forsythe ANOVA test, n = 9, 14 and 6 mice per group; Dunnett’s T3 multiple comparisons test, 0M vs 1M: p = 0.056, 0M vs 2M: p = 0.012; 1M vs 2M: p = 0.236. **Females**: F2,31 = 5.616, p = 0.008, one-way ANOVA, n = 14, 10 and 10 per group; Tukey’s multiple comparisons test, 0M vs 1M: p = 0.011, 0M vs 2M: p = 0.052; 1M vs 2M: p = 0.808). **G.** Anxiety score derived from normalized data from LDT and OFT in males (left panel) and females (right panel) (**Males**: F2,26 = 7.071, p = 0.004, one-way ANOVA, n = 7, 13 and 7 mice per group; Tukey’s multiple comparisons test, 0M vs 1M: p = 0.049, 0M vs 2M: p = 0.386; 1M vs 2M: p = 0.759. **Females**: F2,31 = 11.321, p = <0.001, one-way ANOVA, n = 14, 10 and 10 per group; Tukey’s multiple comparisons test, 0M vs 1M: p = 0.048, 0M vs 2M: p = 0.003; 1M vs 2M: p = 0.246). **H.** After ARS, percentage of time spent in the open arm in a EPM by males (left panel) and females (right panel) (**Males**: F2,28 = 9.287, p = <0.001, one-way ANOVA test, n = 11, 14 and 6 mice per group; Tukey’s multiple comparisons test, 0M vs 1M: p = 0.040, 0M vs 2M: p = <0.001; 1M vs 2M: p = 0.074. **Females**: F2,27 = 5.616, p = 0.007, one-way ANOVA, n = 13, 7 and 10 per group; Tukey’s multiple comparisons test, 0M vs 1M: p = 0.005, 0M vs 2M: p = 0.551; 1M vs 2M: p = 0.065). **I.** After ARS, percentage of time spent in the closed arm in a EPM by males (left panel) and females (right panel) (**Males**: F2,28 = 8.713, p = 0.001, one-way ANOVA test, n = 11, 14 and 6 mice per group; Tukey’s multiple comparisons test, 0M vs 1M: p = 0.033, 0M vs 2M: p = <0.001; 1M vs 2M: p = 0.123. **Females**: F2,28 = 0.235, p = 0.792, one-way ANOVA, n = 13, 8 and 10 per group). Histograms show average ± SEM, * p<0.05, ** p<0.01, ***p<0.001, ns = not significant.

At 3 weeks, immediately after weaning, we found that 1M and 2M mice spent more time in thigmotaxis in the open-field test (OFT) and more time in the dark compartment of the light–dark test (LDT) compared with 0M mice (Fig. 4B, C). This behavioral profile was further captured by a composite anxiety score integrating the two tests (Fig. 4D). Importantly, the total distance travelled was not affected by IUP, indicating that these behavioral differences were not driven by altered locomotor activity (Suppl. Fig. 5A-B). Similarly, at 8 weeks of age, 1M and 2M mice again spent more time in thigmotaxis and in the dark compartment, resulting in a higher composite anxiety score than 0M mice (Fig. 4E-G). Total distance travelled remained unchanged across IUP groups, further supporting the absence of locomotor confounds (Suppl. Fig. 5C and D).

Anxiety reduces resilience to stress, subjecting anxious individuals to an increased risk of depressive-like symptoms after stress exposure. To assess stress reactivity, mice were subjected to acute restraint stress (ARS) for six hours, followed by testing in the elevated plus maze (EPM) the following day. After ARS, 1M and 2M males spent less time in the open arms and risk zone, and more time in the closed arms, compared with 0M males. In females, ARS-induced EPM alterations were less consistent across parameters, suggesting that IUP-related stress reactivity was more robust in males (Fig. 4H and I; Suppl. Fig. 5E). Because postnatal maternal care can profoundly influence offspring emotional development^11, 31–34^, we monitored maternal behavior in the foster dams. We found no IUP-dependent differences in the time dams spent in the nest, suggesting that the observed behavioral effects were unlikely to be explained by gross differences in this measure of maternal care (Suppl. Fig. 5F). Taken together, these findings indicate that natural prenatal variation associated with IUP has long-lasting effects on anxiety-like behavior and shapes stress reactivity in a sex-dependent manner.

### Adult hippocampal neurogenesis is modulated by intra-uterine position

To evaluate the potential impact of IUP on adult hippocampal neurogenesis and its relationship with anxiety-related traits, we quantified hippocampal morphology and neurogenesis in the dentate gyrus (Fig. 5). Following their behavioral assessment, mice were sacrificed at 10 weeks of age, and the volumes of the hippocampus and DG were quantified. In contrast with observations at GD 18.5, 10-weeks-old 1M and 2M mice exhibited reduced hippocampal and DG volumes compared with 0M mice of both sexes (Fig. 5A-D). Notably, this effect appeared specific to the hippocampus and DG, as volumes of the hypothalamus and amygdala were not significantly altered by IUP in either sex (Suppl. Fig. 6C-F). IUP did not affect male body weight, whereas 2M females displayed slightly increased body weight compared with 0M females (Suppl. Fig. 6A). Moreover, both male and female 2M mice showed increased AGD (Suppl. Fig. 6B), in line with previous findings^15^. Together, these anatomical observations indicate that IUP is associated with long-lasting morphological differences extending beyond the prenatal period.

**Figure 5.**
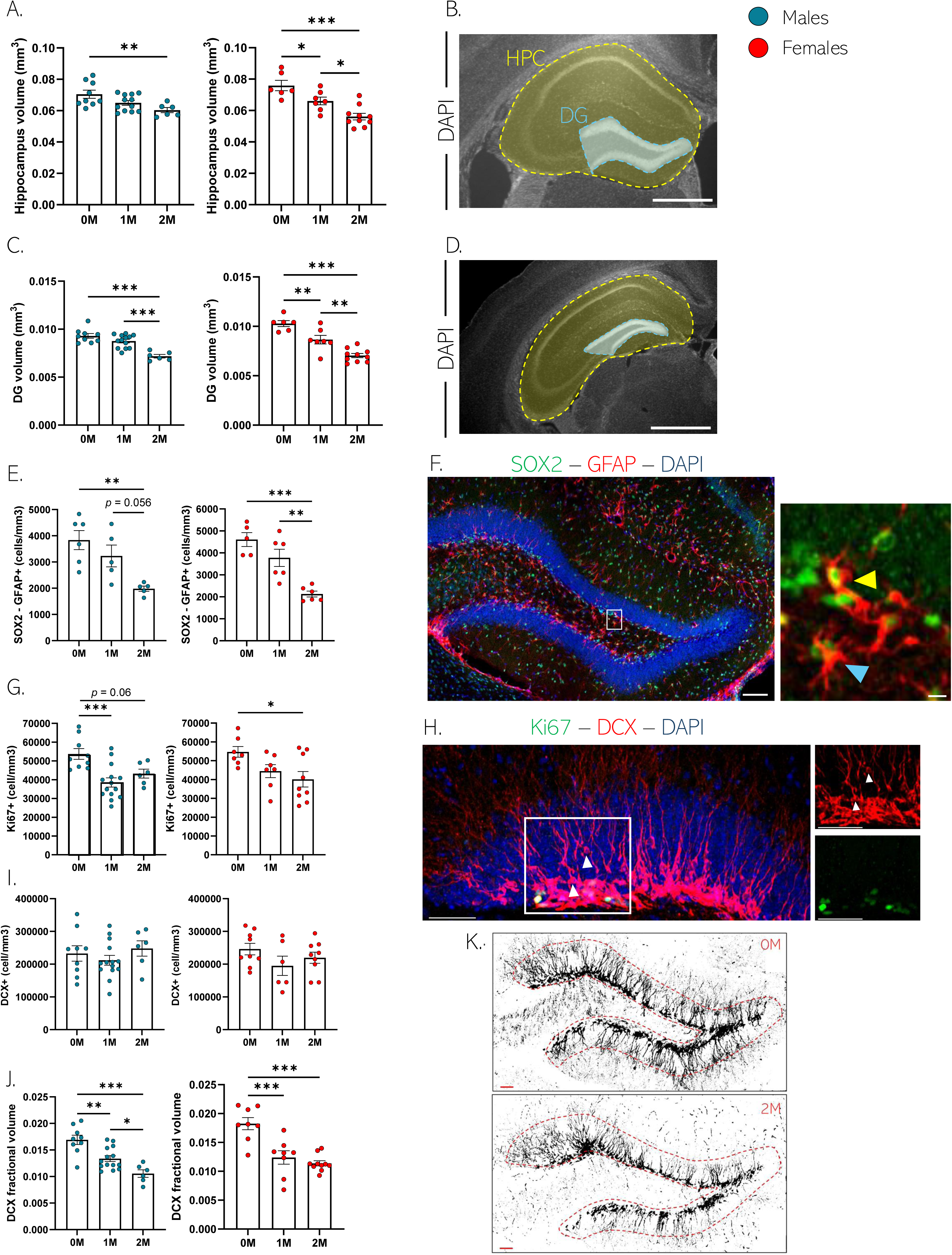
IUP influences hippocampal volume and AHN: **A.** Hippocampal volume in 10-week-old mice, males (left panel) and females (right panel) (**Males**: F2,24 = 5.374, p = 0.012, one-way ANOVA, n = 9, 12 and 6 mice per group; Tukey’s multiple comparisons test, 0M vs 1M: p = 0.114, 0M vs 2M: p = 0.010; 1M vs 2M: p = 0.285. **Females**: F2,20 = 15.56, p = <0.001, one-way ANOVA, n = 6, 7 and 10 per group; Tukey’s multiple comparisons test, 0M vs 1M: p = 0.048, 0M vs 2M: p = <0.001; 1M vs 2M: p = 0.023). **B.** Representative confocal microscopy image of the dorsal hippocampus (highlighted in yellow) and dentate gyrus (highlighted in cyan); immunostained for DAPI (scale bar 1000µm). **C.** Dentate gyrus volume in 10-week-old mice, males (left panel) and females (right panel) (**Males**: F2,25 = 19.24, p = <0.001, one-way ANOVA, n = 9, 13 and 6 mice per group; Tukey’s multiple comparisons test, 0M vs 1M: p = 0.161, 0M vs 2M: p = <0.001; 1M vs 2M: p = <0.001. **Females**: F2,20 = 27.83, p = <0.001, one-way ANOVA, n = 6, 7 and 10 per group; Tukey’s multiple comparisons test, 0M vs 1M: p = 0.007, 0M vs 2M: p = <0.001; 1M vs 2M: p = 0.003). **D.** Representative confocal microscopy image of the ventral hippocampus (highlighted in yellow) and dentate gyrus (highlighted in cyan); immunostained for DAPI (scale bar 1000µm). **E.** Quantification of SOX2-GFAP^+^ cells in males (left panel) and females (right panel) (**Males**: F2,9.46 = 10.58, p = 0.004, Brown-Forsythe ANOVA test, n = 6, 5 and 5 mice per group; Dunnett’s T3 multiple comparisons test, 0M vs 1M: p = 0.555, 0M vs 2M: p = 0.006; 1M vs 2M: p = 0.056. **Females**: F2,10.04 = 17.55, p = <0.001, Brown-Forsythe ANOVA test, n = 5, 6 and 6 per group; Dunnett’s T3 multiple comparisons test, 0M vs 1M: p = 0.325, 0M vs 2M: p = 0.002; 1M vs 2M: p = 0.020). **F.** Confocal maximal projection of DG slices immunostained for DAPI (in blue) SOX2 (in green) and GFAP (in red, blue arrows). Stem cells (yellow arrows) are highlighted by the colocalization of SOX2 and GFAP (scale bar 100µm and 10µm in the inset). **G.** Quantification of Ki-67^+^ cells in males (left panel) and females (right panel) (**Males**: F2,26 = 8.931, p = 0.001, one-way ANOVA test, n = 9, 14 and 6 mice per group; Tukey’s multiple comparisons test, 0M vs 1M: p = <0.001, 0M vs 2M: p = 0.065; 1M vs 2M: p = 0.497. **Females**: F2,20 = 4.170, p = 0.031, one-way ANOVA test, n = 7, 7 and 9 per group; Tukey’s multiple comparisons test, 0M vs 1M: p = 0.168, 0M vs 2M: p = 0.025; 1M vs 2M: p = 0.676). **H.** Confocal maximal projection of DG slices immunostained for DAPI (blue) Ki-67 (green) and DCX (red, white arrows) (scale bar 100µm and 100µm in the inset). **I.** Quantification of DCX^+^ cells in males (left panel) and females (right panel) (**Males**: F2,26 = 0.815, p = 0.454, one-way ANOVA test, n = 9, 14 and 6 mice per group. **Females**: F2,21 = 1.470, p = 0.253, one-way ANOVA test, n = 9, 6 and 9 per group). **J.** DCX fractional volume in the dentate gyrus of the hippocampus of males (left panel) and females (right panel) (**Males**: F2,26 = 15.73, p = <0.001, one-way ANOVA test, n = 9, 14 and 6 mice per group; Tukey’s multiple comparisons test, 0M vs 1M: p = 0.002, 0M vs 2M: p = <0.001; 1M vs 2M: p = 0.037. **Females**: F2,23 = 17.29, p = <0.001, one-way ANOVA test, n = 8, 8 and 10 per group; Tukey’s multiple comparisons test, 0M vs 1M: p = <0.001, 0M vs 2M: p = <0.001; 1M vs 2M: p = 0.689). **K.** Binary images showing the DCX immunolabeled cells in GCL of DG slices of 0M (top panel) and 2M (bottom panel) individuals (scale bar 50µm). Histograms show average ± SEM, * p<0.05, ** p<0.01, ns = not significant.

We then assessed AHN by quantifying radial glia-like cells (RGLs; SOX2^+^ - GFAP^+^) in the subgranular zone and measured cell proliferation in the DG. Both 1M and 2M mice showed reduced RGL cell density and lower proliferation in the subgranular zone, as compared to 0M individuals (Fig. 5E-H). Although the density of immature neurons, identified as DCX^+^ cells, was not significantly affected by IUP (Fig. 5I), markers of immature neuron maturation appeared reduced. This was indicated by lower DCX fractional volume (the fraction of DCX^+^ voxels per region of interest), reduced DCX signal intensity, and fewer dendritic intersections in the upper molecular layer (Fig. 5J, K; Suppl. Fig. 6G-J). Collectively, these results suggest that IUP is associated with persistent differences in adult hippocampal architecture and affects multiple stages of adult hippocampal neurogenesis including stem cell maintenance, proliferation, and neuronal maturation.

### Intra-uterine position alters brain steroid levels

Steroid hormones, including glucocorticoids, testosterone, and progesterone play pivotal roles in shaping brain architecture, by intricately regulating cell proliferation and survival in the hippocampus^35^. To determine whether IUP is associated with altered endocrine states that may contribute to changes in hippocampal neurogenesis, we quantified hormone concentrations at GD 18.5 and 10 weeks of age (Fig. 6A). At GD 18.5, IUP exerted sex-specific effects on brain steroid levels. Specifically, 1M and 2M males exhibited elevated testosterone and progesterone compared with 0M males whereas corticosterone levels were not significantly affected by IUP (Fig. 6B-D, left panels). In contrast, 1M and 2M females showed reduced corticosterone and progesterone levels, while testosterone was undetectable across all females (Fig. 6B-D, right panels). In adulthood, IUP-dependent differences in brain steroid levels persisted, although with a distinct sex-and hormone-specific profile. In males, 2M individuals displayed increased corticosterone levels, whereas testosterone and progesterone were not significantly altered by IUP (Fig. 6E-G, left panels). In females, both 1M and 2M mice showed higher corticosterone and testosterone levels compared with 0M females, while progesterone levels remained comparable across IUP groups (Fig. 6E-G, right panels).

**Figure 6.**
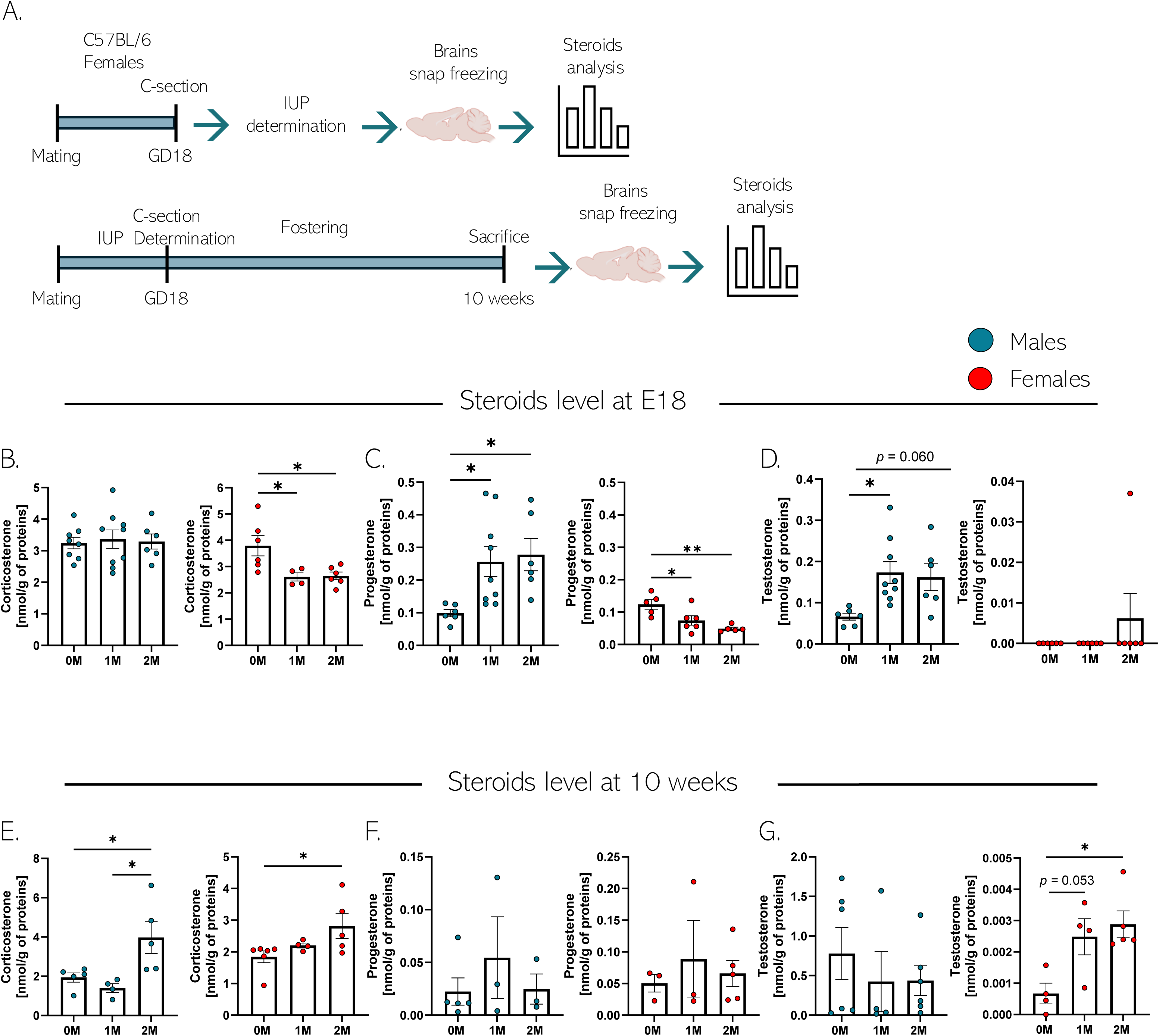
IUP influences brain steroids in embryos and adults: **A.** Schematic representation of the experimental design for hormonal assessment in the brains of embryos (top) and adults (bottom). **B.** Corticosterone levels in the brains of males (left panel) and females (right panel) embryos at GD 18.5 (**Males**: F2,20 = 0.066, p = 0.936, one-way ANOVA test, n = 8, 9 and 6 mice per group. **Females**: F2,13 = 6.123, p = 0.013, one-way ANOVA test, n = 6, 4 and 6 per group; Tukey’s multiple comparisons test, 0M vs 1M: p = 0.034, 0M vs 2M: p = 0.022; 1M vs 2M: p = 0.995). **C**. Progesterone levels in the brains of males (left panel) and females (right panel) embryos at GD 18.5 (**Males**: F2,18 = 4.741, p = 0.022, one-way ANOVA test, n = 6, 9 and 6 mice per group; Tukey’s multiple comparisons test, 0M vs 1M: p = 0.041, 0M vs 2M: p = 0.033; 1M vs 2M: p = 0.930. **Females**: F2,13 = 8.626, p = 0.004, one-way ANOVA test, n = 5, 6 and 5 per group; Tukey’s multiple comparisons test, 0M vs 1M: p = 0.036, 0M vs 2M: p = 0.004; 1M vs 2M: p = 0.354). **D.** Testosterone levels in the brains of males (left panel) and females (right panel) embryos at GD 18.5 (**Males**: F2,18 = 4.991, p = 0.019, one-way ANOVA test, n = 6, 9 and 6 mice per group; Tukey’s multiple comparisons test, 0M vs 1M: p = 0.020, 0M vs 2M: p = 0.060; 1M vs 2M: p = 0.946. **Females**: in females, testosterone was not detected). **E.** Corticosterone levels in the brains of males (left panel) and females (right panel) at 10 weeks of age (**Males**: F2,11 = 6.428, p = 0.014, one-way ANOVA test, n = 5, 4 and 5 mice per group; Tukey’s multiple comparisons test, 0M vs 1M: p = 0.773, 0M vs 2M: p = 0.044; 1M vs 2M: p = 0.018. **Females**: F2,12 = 4.414, p = 0.037, one-way ANOVA test, n = 6, 4 and 5 per group; Tukey’s multiple comparisons test, 0M vs 1M: p = 0.607, 0M vs 2M: p = 0.030; 1M vs 2M: p = 0.234). **F.** Progesterone levels in the brains of males (left panel) and females (right panel) at 10 weeks (**Males**: F2,8 = 0.634, p = 0.555, one-way ANOVA test, n = 5, 3 and 3 mice per group. **Females**: F2,8 = 0.276, p = 0.765, one-way ANOVA test, n = 3, 3 and 5 per group). **G.** Testosterone levels in the brains of males (left panel) and females (right panel) at 10 weeks (**Males**: F2,13 = 0.490, p = 0.623, one-way ANOVA test, n = 6, 4 and 6 mice per group. **Females**: F2,10 = 6.557, p = 0.015, one-way ANOVA test, n = 4, 4 and 5 per group; Tukey’s multiple comparisons test, 0M vs 1M: p = 0.053, 0M vs 2M: p = 0.015; 1M vs 2M: p = 0.881). Histograms show average ± SEM, * p<0.05, ns = not significant.

Overall, these findings indicate that IUP is associated with long-lasting changes in the brain steroid milieu from late embryonic development into adulthood. Together with the observed alterations in hippocampal development and neurogenesis, these endocrine differences support the hypothesis that IUP-dependent variation in prenatal hormone exposure may contribute to persistent changes in brain structure and emotional behavior.

### Prenatal androgen signaling modulates embryonic hippocampal development and neurogenesis

To directly explore the role of steroid hormones on brain anatomy and hippocampal neurogenesis, we employed a parallel approach combining in vitro and in vivo techniques. We first asked which of the steroid hormones altered by IUP in the brain could directly influence neural stem/progenitor cell proliferation. To this end, purified adult hippocampal neural stem/progenitor cells (aNPCs) were exposed for 24 hours to corticosterone, testosterone, or progesterone, and proliferation was assessed by BrdU incorporation (Fig. 7A). Testosterone significantly increased aNPC proliferation compared with control conditions (Fig. 7B and C). In line with previous findings^36^, corticosterone reduced proliferation, whereas progesterone had no significant effect on aNPC proliferation rate. Based on these *in vitro* findings, we hypothesized that altering androgen signaling during development could resemble IUP-related changes. We therefore targeted the androgen receptor (AR) pathway *in vivo* by administering either the AR agonist dihydrotestosterone (DHT), the AR antagonist Flutamide (Flut), or vehicle solution (Veh) to pregnant dams from GD 15 to 17 (Fig. 7D). At GD 18.5, embryos were collected, and brains were analyzed to determine whether prenatal androgen manipulation affected overall embryonic development and neurogenesis within DG. Because DHT increased brain weight in males (Suppl. Fig. 7B), male anatomical measurements were normalized to brain weight. Body weight was not affected by treatment in either sex (Suppl. Fig. 7A).

**Figure 7.**
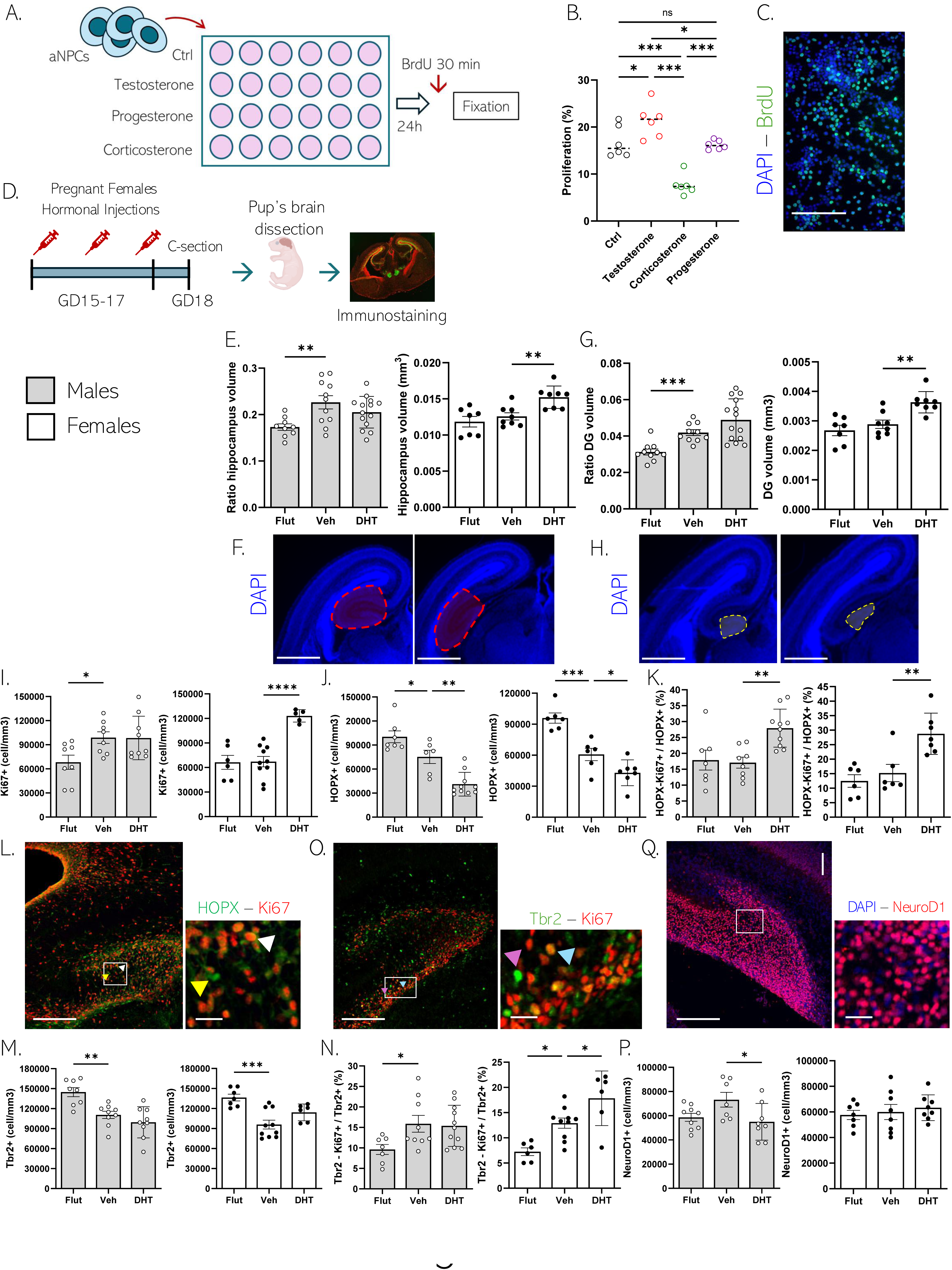
Effects of steroid treatment on aNPC and AHN: **A.** Experimental design for the aNPCs hormonal treatment. **B.** Proportion of aNPC that incorporated BrdU after exposure to steroids for 24 hours under coating conditions (F3,20 = 26.08, p < 0.001, one-way ANOVA; Tukey’s multiple comparisons with the mean of CTRL group, 6 wells per condition, 3 independent replicates). **C.** Representative confocal micrograph of aNPC immunostained for BrdU and DAPI (Scale bar: 100µm). **D.** Schematic representation of the experimental design for assessing the effects of hormonal treatment on brain development and hippocampal neurogenesis. **E.** Analysis of hippocampal volumes in males (left panel) and females (right panel) (**Males**: F2,31 = 5.293, p = 0.010, one-way ANOVA test, n = 9, 11 and 14 mice per group, Dunnett’s T3 multiple comparisons test with the mean of Veh group: Veh vs Flut: p = 0.005, Veh vs DHT: p = 0.266. **Females**: F2,20 = 9.194, p = 0.001, one-way ANOVA test, n = 7, 8 and 8 mice per group, Dunnett’s T3 multiple comparisons test with the mean of Veh group: Veh vs Flut: p = 0.597, Veh vs DHT: p = 0.007). **F.** Representative confocal microscopy image of the dorsal hippocampus (left panel) and ventral hippocampus (right panel) (highlighted in red) immunostained for DAPI (scale bars 1000µm). **G.** Analysis of DG volumes in males (left panel) and females (right panel) (**Males**: F2,24.22 = 15.82, p = <0.0001, Brown-Forsythe ANOVA test, n = 10, 10 and 14 mice per group, Dunnett’s T3 multiple comparisons test with the mean of Veh group: Veh vs Flut: p = 0.0004, Veh vs DHT: p = 0.111. **Females**: F2,20 = 11.57, p = 0.0005, one-way ANOVA test, n = 7, 8 and 8 mice per group, Dunnett’s T3 multiple comparisons test with the mean of Veh group: Veh vs Flut: p = 0.520, Veh vs DHT: p = 0.003). **H.** Representative confocal microscopy image of the dorsal DG (left panel) and ventral DG (right panel) (highlighted in yellow) immunostained for DAPI (scale bar 1000µm). **I.** Quantification of Ki-67^+^ cells in males (left panel) and females (right panel) (**Males**: F2,22 = 4.187, p = 0.029, one-way ANOVA test, n = 8, 8 and 9 mice per group, Dunnett’s T3 multiple comparisons test with the mean of Veh group: Veh vs Flut: p = 0.0373, Veh vs DHT: p = 0.999. **Females**: F2,18 = 17.97, p = <0.0001, one-way ANOVA test, n = 6, 10 and 5 mice per group, Dunnett’s T3 multiple comparisons test with the mean of Veh group: Veh vs Flut: p = 0.9945, Veh vs DHT: p = <0.0001). **J.** Quantification of HOPX^+^ cells in males (left panel) and females (right panel) (**Males**: F2,20 = 23.51, p = <0.0001, one-way ANOVA test, n = 7, 6 and 10 mice per group, Dunnett’s T3 multiple comparisons test with the mean of Veh group: Veh vs Flut: p = 0.0275, Veh vs DHT: p = 0.0023. **Females**: F2,18 = 17.97, p = <0.0001, one-way ANOVA test, n = 6, 10 and 5 mice per group, Dunnett’s T3 multiple comparisons test with the mean of Veh group: Veh vs Flut: p = 0.0348, Veh vs DHT: p = 0.0026). **K.** Percentage of HOPX^+^ - Ki-67^+^ cells over the total number of HOPX^+^ cells in males (left panel) and females (right panel) (**Males**: F2,22 = 8.056, p = 0.0024, one-way ANOVA test, n = 7, 8 and 10 mice per group, Dunnett’s T3 multiple comparisons test with the mean of Veh group: Veh vs Flut: p = 0.9576, Veh vs DHT: p = 0.0032. **Females**: F2,16 = 11.31, p = 0.0009, one-way ANOVA test, n = 6, 6 and 7 mice per group, Dunnett’s T3 multiple comparisons test with the mean of Veh group: Veh vs Flut: p = 0.6988, Veh vs DHT: p = 0.0041). **L.** Representative confocal microscopy image of the dentate gyrus immunostained for HOPX and Ki-67 (HOPX highlighted with yellow arrows and HOPX^+^-Ki-67^+^ highlighted with white arrows; scale bar 100µm and 20µm in the inset). **M.** Quantification of Tbr2^+^ cells in males (left panel) and females (right panel) (**Males**: F2,23 = 11.77, p = 0.0003, one-way ANOVA test, n = 8, 9 and 9 mice per group, Dunnett’s T3 multiple comparisons test with the mean of Veh group: Veh vs Flut: p = 0.0034, Veh vs DHT: p = 0.3996. **Females**: F2,20 = 11.50, p = 0.0005, one-way ANOVA test, n = 7, 10 and 6 mice per group, Dunnett’s T3 multiple comparisons test with the mean of Veh group: Veh vs Flut: p = 0.0002, Veh vs DHT: p = 0.0957). **N.** Percentage of Tbr2^+^ - Ki-67^+^ cells over the total amount of Tbr2^+^ cells in males (left panel) and females (right panel) (**Males**: F2,24 = 3.653, p = 0.0412, one-way ANOVA test, n = 7, 9 and 11 mice per group, Dunnett’s T3 multiple comparisons test with the mean of Veh group: Veh vs Flut: p = 0.0399, Veh vs DHT: p = 0.9644. **Females**: F2,19 = 12.56, p = 0.0003, one-way ANOVA test, n = 6, 10 and 6 mice per group, Dunnett’s T3 multiple comparisons test with the mean of Veh group: Veh vs Flut: p = 0.0143, Veh vs DHT: p = 0.0332). **O.** Representative confocal microscopy image of the dentate gyrus immunostained for Tbr2 and Ki-67 (Tbr2 highlighted with pink arrows and Tbr2^+^-Ki-67^+^ highlighted with cyan arrows; scale bar 100µm and 20µm in the inset). **P.** Quantification of NeuroD1^+^ cells in males (left panel) and females (right panel) (**Males**: F2,21 = 3.643, p = 0.0438, one-way ANOVA test, n = 9, 7 and 8 mice per group, Dunnett’s T3 multiple comparisons test with the mean of Veh group: Veh vs Flut: p = 0.0845, Veh vs DHT: p = 0.0325. **Females**: F2,20 = 0.378, p = 0.6899, one-way ANOVA test, n = 7, 8 and 8 mice per group, Dunnett’s T3 multiple comparisons test with the mean of Veh group: Veh vs Flut: p = 0.9112, Veh vs DHT: p = 0.8268). **Q.** Representative confocal microscopy image of the dentate gyrus immunostained for NeuroD1 (in red) and DAPI cells (in blue) (scale bar 100µm and 20µm in the inset). Histograms show average ± SEM, * p<0.032, **p<0.0021, ***p<0.0002, ****p<0.0001, ns = not significant.

Prenatal androgen manipulation produced sex-specific effects on embryonic brain development. In females, Flutamide selectively reduced hypothalamic volume without affecting other regions (Suppl. Fig. 7C–E, right panels) whereas DHT increased hippocampal and DG volumes compared with vehicle-treated controls (Fig. 7E-H, right panels). In males, Flutamide reduced cortical thickness and decreased hypothalamic, hippocampal, and DG volumes compared with vehicle-treated embryos, while amygdala volume was not significantly affected (Fig. 7E-H, left panels; Suppl. Fig. 7C-E, left panels). In contrast, DHT did not further increase regional brain volumes in males compared with vehicle-treated controls.

We next assessed whether prenatal androgen manipulation could recapitulate IUP-related changes in embryonic DG neurogenesis. In females, DHT increased overall cell proliferation and enhanced proliferation within neural stem cell and progenitor populations (Fig. 7I, K, L, N, and O, right panels). This was accompanied by a reduction in the HOPX^+^ neural stem cell population (Fig. 7J, right panel). Conversely, Flutamide increased the number of HOPX^+^ and Tbr2^+^ cells (Fig. 7J and M, right panels) and reduced progenitor proliferation (Fig. 7N, right panel). In males, DHT selectively increased proliferation within the neural stem cell population without affecting overall proliferation (Fig. 7I, K and L, left panels). Flutamide, instead, was associated with reduced neurogenic activity, including fewer proliferating cells and lower Tbr2^+^ progenitor proliferation compared with vehicle-treated embryos (Fig. 7I and N, left panels). DHT modestly reduced the number of NeuroD1^+^ immature neurons in males, whereas no significant effect was observed in females. (Fig. 7P and Q).

In summary, prenatal androgen manipulation exerted sex-specific effects on embryonic hippocampal development and DG neurogenesis. In females, DHT promoted hippocampal and DG growth and enhanced proliferation across neurogenic populations, partially resembling the effects observed in embryos positioned adjacent to males in utero. In males, Flutamide reduced hippocampal development and neurogenic activity, whereas DHT did not further enhance most developmental outcomes. Together, these findings support a role for androgen signaling in embryonic hippocampal neurogenesis and raise the possibility that IUP-dependent variation in prenatal androgen exposure contributes to the developmental effects of intra-uterine position.

## Discussion

In this study, we investigated the IUP as a potential non-genomic source of interindividual variability in trait anxiety and stress resilience. We found that embryos positioned adjacent to male littermates (i.e. 1M and 2M) displayed increased volume in several brain areas of the limbic system, with a large effect size on the dentate gyrus. In the hippocampus, gene expression analyses revealed a distinct transcriptomic profile in stem cells indicating reduced stemness and early activation of a neurogenic program involving androgen-dependent transcriptional regulation. The brains of 1M/2M embryos displayed a distinct hormonal profile as compared to 0M mice and manipulation of androgen signaling partially recapitulated several IUP-associated morphological and cellular phenotypes. Later in life, 1M/2M mice displayed reduced hippocampal volume and AHN, and increased anxiety-like behavior. Together, these findings indicate that IUP initiates a cascade of developmental events that contribute to interindividual differences in hippocampal plasticity, emotional behavior and stress vulnerability in adulthood.

A central implication of these findings is that behavioral and anatomical variability among inbred animals housed in standardized conditions should not be viewed only as experimental noise but may reflect subtle variations occurring during development. IUP provides a particularly striking example of this principle by generating distinct microenvironments among embryos within the same litter and can be considered a form of prenatal non-shared environment: while siblings share the same mother and postnatal period, few experience the same local hormonal milieu during gestation. This observation is consistent with the broader concept that non-shared environmental factors contribute substantially to behavioral and brain structural differences among genetically related individuals^8–10, 37, 38^. Our data extend this framework by suggesting that such non-shared influences may begin before birth, through the spatial organization of embryos within the uterus. Mechanistically, our findings support a model in which IUP-associated steroid exposure alters the developmental dynamics of hippocampal neural stem and progenitor cells. During late embryonic development, 1M and 2M embryos displayed increased proliferation and changes in neurogenic progression. In adulthood, however, these same IUP groups showed fewer radial glia-like stem cells and reduced proliferative activity in the dentate gyrus. As HOPX^+^ neural progenitors in the developing dentate gyrus transition to HOPX^+^ quiescent neural stem/progenitor cells in the adult^39^, our observations suggest that increased embryonic neurogenesis may occur at the expense of long-term stem cell maintenance. Such a mechanism is consistent with previous studies showing that developmental acceleration of neurogenesis can transiently increase neuronal production while ultimately depleting the progenitor pool and impairing adult neurogenesis later in life^40–42^. In line with this interpretation, our single-cell transcriptomic data further support the idea that IUP is associated with altered neural stem cell states, with 1M and 2M cells showing signatures compatible with increased neurogenic commitment and reduced long-term stemness. Thus, IUP may influence not only the number of developing progenitors, but also their transcriptional identity and lifelong neurogenic potential.

The behaviors observed at PND22-23 and at 8 weeks of age indicate increased anxiety and stress reactivity in 1M and 2M mice. These behavioral findings are consistent with the known role of AHN in affective regulation and stress adaptation. Experimental reduction of AHN has been associated with increased anxiety-like behavior and stress vulnerability, whereas interventions that enhance AHN can promote resilience to anxiety-and depression-like phenotypes^43–46^. In addition, natural variations in AHN inversely correlate with trait anxiety^47, 48^. Thus, by depleting the neural stem cell pool in the dentate gyrus later in life, the vicinity of males during embryonic development may contribute to increased anxiety-related behavior adolescence to young adult mice.

Our hormonal manipulation experiments support androgen signaling as a mechanism contributing to IUP’s effects. These results are consistent with the hypothesis that IUP modifies the embryos’ exposure to androgens^13, 15, 49^ and the expression of masculinized anatomical and behavioral traits^16, 17^. However, we also observed changes in other steroid hormones, including corticosterone and progesterone, suggesting that IUP may influence a broader steroidal microenvironment. This is particularly relevant because glucocorticoids and sex steroids can have distinct, and sometimes opposing, effects on hippocampal neurogenesis^50–52^. Although non-hormonal effects cannot be excluded, the phenotype observed in 1M and 2M animals may therefore result from the combined influence of multiple steroid hormones acting during sensitive windows of hippocampal development.

Together, the present study identifies the embryonic environment as an important source of variability that shapes prenatal steroid exposure, hippocampal development, adult neurogenesis, and eventually, anxiety-related behavior in mice. These results are relevant to brain health in the context of twin pregnancies^53^ or elevated androgens in pregnant mothers^29^.

## Acknowledgements

The authors wish to thank, Fulvio Magara, Morgane Tuscello, Fanny Thevenaz and the group of the Professor Jose Luis Trejo for help and advice with the behavioral experiments. We would also like to thank the caretakers from the Centre d’Etude Comportementale (CEC). snRNA sequencing was performed at the genomics facility of the University of Lausanne. Hormones measurements was performed at the metabolomics platform of the University of Lausanne. This project was funded by the Swiss National Science Foundation (Grant Number 310030_201015).

## Author contributions

FG, TL, and NT designed research. FG, MV, MA, PvG and TS performed research. FG, MV, MA, PvG, TS, HG, JI, TB, AH and NT analyzed the data. FG and NT wrote the paper. TL and NT supervised the research. NT obtained the funding.

## Material and methods

### Animals

All experiments were performed with the approval of the Cantonal Veterinary Authorities (Vaud, Switzerland) and carried out in accordance with the European Communities Council Directive of 24 November 1986 (86/609EEC). All experiments were performed on C57BL/6J male and female mice obtained from Janvier Laboratories (France). Mice were weighed weekly to track their health status and pregnancy. Mice were maintained under standard housing conditions on corncob litter in a temperature-(23 ± 1°C) and humidity-(40%) controlled animal room with a 12 h light/dark cycle (8h00–20h00), with unlimited access to food and water. All tests were conducted during the light period.

#### Maternal behavior monitoring

Starting on PND1, maternal behavior was monitored twice daily, at 09:00 and 18:00, for two weeks during the light phase, when nursing behavior is most frequently observed. Using an adapted snapshot-based protocol^54^, each cage was observed for 5 seconds every 4 minutes, with each session comprising 10 snapshots per cage, resulting in a total of 280 observations per dam across the monitoring period.

For each observation, foster dams were scored as either being in the nest or outside the nest. The total time spent in the nest was used as a proxy measure of maternal care. This approach was based on our previous findings showing that time spent in the nest correlates with more detailed maternal care parameters, including licking/grooming and arched-back nursing^11^. Time spent in the nest was then compared across foster dams assigned to the different IUP groups to assess whether gross differences in maternal care could account for the behavioral outcomes observed in the offspring.

#### Acute restraint stress (ARS)

The acute restraint stress protocol is based on a previously described protocol^55^. Animals were introduced headfirst into 50 ml Falcon tubes (11.5 cm in length; diameter of 3 cm) from which the cap was removed, and the bottom was perforated with four 0.4 cm holes to enable breathing. Tissue was added at the caudal extremity to adjust the physical constraint to the mouse size and to allow the tail to expand out of the tube. The mice were subjected to this restrained environment for 6 consecutive hours. After one 6-hours-restraining period, mice were transferred into their home-cage followed by an EPM test 24 hours later, to assess anxiety-like behavior.

### IUP determination

Two virgin C57BL/6J females were housed with one C57BL/6J male in the male’s home cage for mating. The following morning, females were checked for the presence of a vaginal plug, and this day was designated as gestational day 0.5 (GD0.5). Males were then removed, and baseline body weight was recorded for subsequent monitoring of pregnancy progression and general health. Pregnancy was confirmed on GD11 by an increase in body weight of at least 30% compared with GD1, together with an observable increase in abdominal/hip circumference. On GD 18.5, corresponding to the expected day of parturition, pregnant dams were euthanized by cervical dislocation, and the uterus was rapidly exposed. Embryos were carefully dissected from the amniotic sacs one by one, preserving their relative order within the uterine horns. For each embryo, sex was determined based on anogenital distance (AGD), measured from the center of the genital papilla to the proximal end of the anus, as described by Manno^56^. AGD is a reliable marker of fetal sex, with males displaying a longer AGD than females^15^. The presence of scrotal pigmentation was used as an additional criterion to identify male embryos, whereas female embryos lacked pigmentation in the anogenital region^57^. After sex determination, each embryo was weighed.

IUP was assigned according to the sex of the immediately adjacent embryos within the uterine horn. Embryos with no adjacent male were classified as 0M, embryos with one adjacent male as 1M, and embryos with two adjacent males as 2M. Embryos located at the extremities of the uterine horns, which had only one direct neighbor, were classified according to the sex of that neighbor: terminal embryos adjacent to a female were classified as 0M, whereas terminal embryos adjacent to a male were classified as 1M. The two uterine horns were not treated as separate experimental factors in downstream analyses.

### Fostering

The gestation of C57BL/6J and CD1 foster dam were synchronized so that CD1 dams delivered 1–2 days before C57BL/6J pups were obtained by caesarean section.

On GD18.5, C57BL/6J pregnant dams underwent caesarean section, and pups were collected for IUP determination as described above. After extraction from the amniotic sac, C57BL/6J pups were gently stimulated on a heating pad to promote breathing and were then placed in an incubator pre-heated to 37°C until breathing and body temperature were stabilized. Pups were then grouped according to their IUP classification.

Before fostering, approximately half of the biological CD1 litter was removed, up to a maximum of four CD1 pups with the foster dam. Each CD1 foster dam received a maximum of eight C57BL/6J pups. Each foster dam received C57BL/6J pups from only one IUP group: 0M foster dams received only 0M pups, 1M foster dams received only 1M pups, and 2M foster dams received only 2M pups. Litter size and sex ratio were balanced across foster litters. After transfer, pups were left undisturbed with the CD1 foster dam and the remaining CD1 pups, except for routine husbandry and experimental monitoring.

### Single-nucleus RNA seq

For single-nucleus RNA sequencing, hippocampi were dissected from GD 18.5 embryos following IUP determination. The final dataset included seven embryos: 0M females, n = 1; 0M males, n = 2; 1M females, n = 1; 1M males, n = 1; and 2M males, n = 2. Samples were not pooled at the tissue-dissection level. Nuclei were isolated separately from each embryo using the 10x Genomics Nuclei Isolation Kit, and sample identity was preserved during library preparation and downstream demultiplexing through sample-specific barcoding.

Nuclei were processed using the 10x Genomics Chromium GEM-X Single Cell 5’ v3 Gene Expression platform according to the manufacturer’s instructions. Library preparation and sequencing were performed at the University of Lausanne Genomics Facility, using Illumina NovaSeq 6000. Raw sequencing data were processed using Cell Ranger v.8 with default parameters and aligned to the mouse reference genome (GRCm39-2024-A). Downstream analysis was performed in R using Seurat v5^58^. Only genes expressed in at least three nuclei were retained. Nuclei were filtered to include cells expressing at least 500 unique genes, with less than 2% mitochondrial transcripts and less than 2.5% ribosomal transcripts. Normalization and variance stabilization were performed using SCTransform. Dimensionality reduction was performed using principal component analysis (PCA), and the resulting principal components were used to calculate UMAP coordinates. A K-nearest neighbor graph was generated, and cells were clustered using the Louvain algorithm. Cell types were annotated based on known marker gene expression and with reference to published mouse brain datasets^59^ using Seurat’s FindTransferAnchors and TransferData functions with SCT normalization. Predicted annotations were then manually refined based on cluster-specific marker genes and grouped into major hippocampal and adjacent cortical cell populations.

#### NSC subtype proportion analysis

For each mouse, the proportion of each NSC subpopulation was calculated as the number of cells assigned to that population divided by the total number of NSCs. Differences in cell-type proportions between the 1M/2M and 0M groups were assessed using quasibinomial generalized linear models fitted separately for each population. Models were specified as cbind(n, n_other) ∼ group, where n represents the number of cells assigned to the population of interest and n_other represents all remaining cells within the corresponding denominator population. Quasibinomial models were used to account for overdispersion across biological replicates relative to binomial expectations. P-values were adjusted across tested populations using the Benjamini-Hochberg method.

#### Differential gene expression analysis

Differential gene expression between 1M/2M and 0M groups was performed in Seurat within each cell type. Differential expression was performed using the MAST test with logfc.threshold = 0.25, min.pct = 0.1, and latent variables nCount_RNA, percent.mt and sex. For IUP-specific and sex-specific comparisons, a similar approach was used except for the omission of sex as a latent variable. Genes were considered significant when Benjamini–Hochberg-adjusted p-values were < 0.05 and absolute average log2 fold-change was > 0.25.

#### Gene ontology enrichment analysis

Gene Ontology (GO) enrichment analysis was performed using clusterProfiler in enrichGO^60^. Significant NSC DEGs were used as input gene lists, with all genes detected in NSCs used as the background universe. GO enrichment was performed using org.Mm.eg.db, keyType = “SYMBOL”, ontology set to biological process (ont = “BP”), Benjamini Hochberg adjustment, and pvalueCutoff = 0.05. GO terms with adjusted p-values < 0.05 were retained. For each enriched term, gene ratio, background ratio, fold enrichment, and -log10(adjusted p-value) were calculated. A composite enrichment score was calculated as fold enrichment multiplied by -log10(adjusted p-values) for ranking and visualization. GO annotations were further annotated using GO.db^61^ (DOI: 10.18129/B9.bioc.AnnotationDbi) to retrieve GO terms, ontology labels, definitions, and direct parent terms. Semantic parent terms were used to reduce redundancy and curate higher-order biological categories. GO visualizations included dot plots, semantic parent summaries, and circular pathway representations using GOplot^62^.

#### Slingshot pseudotime trajectory inference

Trajectory inference was performed using Slingshot in a SingleCellExperiment framework^63^. A neural lineage subset, referred to in the analysis scripts as the neu subset, was used for NSC to neuronal trajectory analysis. This subset was reprocessed in Seurat using the RNA assay, SCTransform, PCA with 50 principal components, UMAP using dimensions 1:30, nearest-neighbor graph construction using dimensions 1:30, and clustering using FindClusters (resolution = 0.4). The Seurat object was converted to a SingleCellExperiment object, and Slingshot was run on PCA coordinates using curated cluster labels. We performed Slingshot trajectory inference using Hopx^+^/Mki67^+^ RGL as the biologically defined root cluster. Terminal states were first inferred without specifying endpoints to assess whether the data topology supported the expected neurogenic branches. In a subsequent hypothesis-guided analysis, CA1, CA3, and DG were used as terminal states to estimate lineage-specific pseudotime for downstream tradeSeq testing. Slingshot was run with stretch = 0. Pseudotime values were extracted from Slingshot^64^ lineage assignments, and when multiple lineage-specific pseudotime values were present for a cell, average pseudotime was calculated across available lineages. Cells without valid pseudotime values were excluded from pseudotime-dependent summaries. Slingshot trajectories were visualized in PCA space for interpretation of NSC to neuronal progression.

#### TradeSeq trajectory dependent gene expression modeling

Trajectory associated gene expression was modeled using tradeSeq^65^. Raw RNA counts were extracted from the trajectory subset and filtered to retain genes expressed in at least 10 cells. Generalized additive models were fitted using fitGAM with the Slingshot object, filtered count matrix, nknots = 6, and conditions = group, where group encoded 0M versus 1M/2M status. The following tradeSeq tests were applied:

- startVsEndTest(global = TRUE, lineages = FALSE) to identify genes changing between early and late pseudotime states.
- conditionTest(l2fc = log2(2)) to identify genes showing condition-associated expression differences along pseudotime.
- patternTest(l2fc = log2(2)) to identify genes with altered expression dynamics or trajectory patterns between groups.

P-values from tradeSeq tests were adjusted using the Benjamini Hochberg method. Representative genes were visualized using fitted smoother curves and pseudotime-binned expression summaries. Early to late differentiation markers such as *Mki67* and *Dcx* were interpreted using startVsEndTest, while group associated trajectory effects such as persistent *Mef2c* upregulation in 1M/2M relative to 0M were interpreted using conditionTest.

#### Transposable element analysis using SoloTE

Transposable element (TE) expression was analyzed using SoloTE^66^ derived count matrices. SoloTE outputs were transferred into the final Seurat object as independent assays, including SoloTE_legacy, SoloTE_subfamily, and SoloTE_locus. Barcode formats were harmonized between merged Seurat cells and SoloTE outputs before assay transfer. TE expression was analyzed at subfamily resolution. Group and cluster associated TE differences were tested within each group or cluster by setting identities to group and applying FindMarkers between the two IUP groups (0M and 1M/2M) or between Neurogenic RGL and other RGL.Gliogenic clusters with min.pct = 0.1, logfc.threshold = 0.25, and MAST testing with latent.vars (nCount_RNA, percent.mt and sex). Cluster wise TE differential analyses were skipped when either comparison group contained fewer than three cells.

#### Data visualization and statistical analysis

All computational analyses were performed in R using Seurat, SingleCellExperiment, Slingshot, tradeSeq, tidyverse^67^, broom^68^, clusterProfiler, GO.db, AnnotationDbi, GOplot, and related visualization packages. Unless otherwise stated, adjusted p-values < 0.05 was considered statistically significant.

### Steroids hormones quantification

Steroid hormone levels were quantified in embryonic and adult brain tissue by LC-MS/MS. A panel of twenty-four endogenous steroids was targeted for quantification in brain tissue lysates, as described below.

For embryonic samples, whole brains were collected at GD 18.5 immediately after IUP determination and snap-frozen in pre-chilled 70% ethanol before storage at −80°C. For adult samples, mice were sacrificed at 10 weeks of age after completion of the behavioral experiments. One brain hemisphere was randomly selected for steroid hormone quantification, immediately snap-frozen in pre-chilled 70% ethanol, and stored at −80°C until processing. To minimize potential circadian variability in steroid hormone levels, all brain dissections were performed in the morning, between 09:30 and 11:30.

#### Standard solutions

Calibrators (i.e., standard solutions) were prepared in depleted serum samples. A mixture of IS (IS-mix with authentic IS for each steroid) was prepared by spiking different volumes of each IS solution at appropriate concentrations for each IS in MeOH.

#### Sample preparation

Steroid hormones were extracted using Solid phase extraction (SPE) on Oasis Prime HLB 96-well plates (Waters). For each sample, the embryonic and adult brain tissue (∼10mg) was first extracted by the addition of 150 μL of the IS mixture in MeOH using tissue homogenizer or bead beater. Following homogenization, the tissue lysate was diluted by the addition of 500 μL of PBS and 250 μL of 0.1 M ZnSO4, mixed and centrifuged. Following SPE column conditioning, the prepared supernatants of tissue lysates (as described above) were loaded onto the columns, and positive pressure was applied to facilitate sample loading and adsorption. Following washing (with 80% H2O), the elution was carried out with acetonitrile and methanol (500 μL ACN/MeOH 1:1, v/v). Eluates were collected in collection plates and evaporated under the stream of nitrogen and finally reconstituted with 50 μL of MeOH/H2O 1:1 (v/v) solution for LC-MS/MS analysis.

#### LC-MS/MS analysis

Each extract was analyzed by ultra-high performance liquid chromatography (UHPLC) coupled to a Q-TRAP 6500 plus triple quadrupole mass spectrometer (Sciex) operating in positive and in negative ionization mode. Liquid chromatography was performed using a Premier ACQUITY UPLC HSS T3 column (100 × 2.1 mm, 1.8 μm; Waters) set at 50 °C. The mobile phase A was a solution of 0.25 mM of ammonium fluoride in H2O, and the mobile phase B was 0.25 mM ammonium fluoride in MeOH. The gradient elution in positive mode was performed starting with 20% B for the first 0.25 minutes. The percentage for solvent B increased to 46% at 0.5 minutes and held constant until 1 minute. It was then raised to 54% by 1.25 minutes. From 1.25 to 4.75 minutes, the percentage of solvent B decreased to 42%, followed by a further reduction to 35% at 5 minutes. Over the next two minutes, solvent B was reduced to 30% at 7 minutes, and then to 26% at 8.25 minutes. A sharper decrease occurred, bringing solvent B to 10% by 9.5 minutes, and to 1% at 9.75 minutes. The gradient was maintained at 1% until 11.75 minutes, after which it was increased to 20% at 12 minutes, and held constant until 15 minutes. The gradient elution in negative mode began with 20% B for the first 0.25 minutes, after which the percentage of solvent B was increased to 53% by 0.5 minutes and held constant until 1 minute. From 1 to 2 minutes, the solvent B was raised to 68% and maintained at this level for 3 minutes. A gradual increase was then applied, bringing the percentage to 72% by 3.5 minutes, followed by a further increase to 83% by 5 minutes. The percentage for solvent B reached 99% by 5.25 minutes and was held at this level until 12 minutes. The injected volume was 5 μL and the flow rate was set at 450 μL/min. ESI-MS/MS parameters were set as follows: curtain gas (CUR) was set to 20, and the collision gas (CAD) was maintained at a low setting. The ion spray voltage (IS) was 5500 V in positive mode and -4500 V in negative mode. The source temperature (TEM) was adjusted to 400°C for positive mode and 600°C for negative mode. The ion source gases were set as follows: Ion Source Gas 1 (GS1) at 50 and Ion Source Gas 2 (GS2) at 60. The entrance potential (EP) was 10 V for positive mode and -10 V for negative mode. The system operated in Scheduled Multiple Reaction Monitoring (MRM) mode to maximize dwell times and optimize cycle times, with a target scan time of 0.4 seconds per sMRM experiment. The minimum dwell time was set to 10 milliseconds, and the resolution for Q1 and Q3 was configured in unit mode. Ion Spray voltage, collision energy, and MS/MS transitions were optimized by infusing standard solutions of each compound at 100 ng/mL in reconstitution solvent. Data were processed using MultiQuant software (version 3.0.3).

### Cell culture

Adult hippocampal neural stem/progenitor cells (aNPCs) were isolated from the hippocampus of adult Fischer 344 rats and cultured in Dulbecco’s Modified Eagle Medium/Nutrient Mixture F-12 (DMEM/F12) supplemented with B27 and fibroblast growth factor 2 (FGF-2; 20 ng/mL), as previously described^47^. After isolation, cells were expanded, split, cryopreserved, and thawed for experimental use. For steroid exposure experiments, aNPCs were plated at a density of 60,000 cells per well in 500 μL of medium in 24-well plates coated with poly-D-lysine hydrobromide (0.1 mg/mL; Sigma-Aldrich, P6407), poly-L-ornithine, and laminin (20 μg/mL; Life Technologies, 23017015). Twenty-four hours after plating, the medium was removed and replaced with fresh medium containing corticosterone (100 nM; Sigma-Aldrich, 50-22-6), testosterone (20 nM; Sigma-Aldrich, 58-22-0), progesterone (10 nM; Sigma-Aldrich, 57-83-0), or fresh medium alone as control condition. Steroid stock solutions were prepared in ethanol and diluted in culture medium immediately before use. Cells were exposed to steroid hormones for 24 h. Following steroid exposure, 5 μM BrdU was added to the culture medium for 30 min. Cells were then washed twice with pre-warmed fresh medium and fixed with 4% paraformaldehyde for 30 min. Plates were briefly washed with 0.1 M PBS and stored at 4°C until immunocytochemistry. For BrdU immunostaining, DNA was denatured with 2 M HCl for 15 min at 37°C, followed by incubation in 0.1 M borate buffer, pH 8.5, for 15 min. Cells were then incubated for 1 h in blocking solution containing 0.3% Triton X-100 and 10% heat-inactivated horse serum (Gibco, 16050122). Plates were incubated for 24 h at 4°C with mouse monoclonal anti-BrdU antibody (1:250; Abcam), followed by incubation with goat anti-mouse Alexa Fluor 488 secondary antibody (1:300; Invitrogen) for 1 h at room temperature. Nuclei were counterstained with DAPI (1:500) for 10 min. Images were acquired using a Nikon microscope and analyzed automatically with Nikon imaging software by an experimenter blinded to the experimental conditions. For each well, six fields were analyzed, and the percentage of BrdU-positive cells was calculated over the total number of DAPI-positive cells.

### Hormones administration

The steroid hormones selected for this experiment were DHT (Sigma Aldrich, A8380) at a concentration of 2.5mg/kg body weight dissolved in a vehicle solution composed of a 1:1 ratio of corn oil (Sigma Aldrich, C8267) and benzyl benzoate (Sigma Aldrich, B6630). Flutamide (USP, 1285851) was prepared at a concentration of 120mg/kg body weight dissolved in the same vehicle. The hormones doses were adapted from Zheng and Cohn, 2011. Vehicle was administered as the control. Each pregnant female received one subcutaneous injection of the assigned treatment (vehicle, DHT, or Flutamide) daily from GD15 to GD17, around the same time each day. The total injected volume was 0.2 mL for the vehicle and 0.15 mL for both Flutamide and DHT. Special care was taken during injections to ensure no pressure was applied to the uterus during restraint, minimizing harm to the mice.

### Anxiety tests

#### Light - Dark test (LDT)

As previously described^69^, the apparatus used for the LDT consisted of a white wooden box with two compartments. One was a square compartment without a lid, serving as the light side (40 x 40 cm), while the other was a smaller rectangular compartment with a lid, creating the dark side (20 x 40 cm). These two compartments were connected by a 5 x 5 cm door, and the entire apparatus had a height of 30 cm. The center of the lit compartment maintained a stable luminosity of 400 lux, while the dark compartment remained without any light source. Mice were introduced into the apparatus in the light side, facing the door, and allowed to explore for a duration of 5 minutes. The mice’s movements were tracked and recorded using the ANY-maze software. In this test, anxiety-like behavior was evaluated based on the time spent in the dark compartment.

#### Open field test (OFT)

The OFT was conducted as previously described in^70^, in a rectangular arena (50 × 50 × 40 cm^3^) illuminated with dimmed light (25 lux). Mice were introduced near the wall of the arena and allowed to explore for 10 min. Analyses were performed using ANY-maze tracking software by drawing a virtual zone (15 × 15 cm^2^) in the center of the arena defined as the anxiogenic area. Several parameters were analyzed, including the total distance travelled and the time spent in the different zones.

#### Elevated plus maze (EPM)

The test was conducted as previously described in^70^. Briefly, the animals were placed in a maze made from black PVC with a white floor. The EPM consists of 4 arms (30 × 5 cm) arranged in a plus shape, with walls on two opposing arms, while the other two arms are exposed to the height of the apparatus (65 cm). The arms converge at the center, forming a small, squared platform (5 × 5 cm). Throughout the tests, the light conditions were maintained stable. The open arms and the center of the maze had a luminous flux of 12–15 lux, while the closed arms had reduced light intensity with 5 lux. The animals were gently introduced into the maze, facing the wall at the end of the closed arms, and were allowed to freely move within the maze for a duration of 5 min. Video recordings of the mice’s behavior were captured from above the arena, and tracking analyses were performed using the ANY-maze software to determine the time spent in both the open and closed arms.

### Anxiety score

The anxiety scores encompassed several anxiety tests to get a general profile of anxiety as previously described^70–72^. The score is derived from the normalization of the values for the combination of individual anxiety tests (time spent in the dark chamber during LDT and time spent in thigmotaxis in an OFT). The normalization involved adjusting each animal’s value by subtracting the minimum value of the entire population and then dividing this result by the difference between the maximum and minimum values of the entire population: (x – min value) / (max value – min value). This method generates scores distributed on a scale from 0 to 1, with a score of 1 indicating high anxiety.

### Samples collection and preparation

#### GD 18.5 brain preparation

After IUP determination embryo’s body weight was recorded. Embryos were decapitated; the brain was dissected and weighed. Immediately after, the brain was stored in 2mL of 4% PFA (Sigma Aldrich, 158127) in 15mL tubes for post-fixation and then stored at -20°C. 24 hours later, the brains were carefully removed from the PFA and transferred into gradients of sucrose (15% and 30%) to dehydrate the brain for optimal freezing. They were then placed into OCT embedding medium (Leica Microsystems, 14020108926) and snap frozen using dry ice and stored at -80°C until slicing. The brains were transferred to -20°C one day prior to cutting. 35um thick coronal frozen sections of the dorsal and ventral hippocampus were obtained using a cryostat (Leica Microsystems CM3050S) at a controlled temperature of -22°C. The slices were directly collected onto gelatine coated (Superfrost PLUS slides Epredia, (J1810AMNZ)) and stored in slide holders containing ∼80mL of PBS 1x (sufficient to cover all slices) at RT until immunohistochemistry staining. In a second cohort, to assess steroid hormone levels in the brain, dissected brains were immediately submerged in 70% ethanol pre-chilled to –60/–70 °C for snap-freezing. Samples were then stored at –80 °C until further processing.

#### 10-week-old adult brains

After all the behavioral experiments the mice were sacrificed with a lethal injection of pentobarbital (10 mL/kg, Sigma-Aldrich, Switzerland) and transcardially perfused with saline solution (NaCl 0.9%). Mice were weighed and AGD was measured. Brains were removed and half sectioned to then perform immunohistochemistry and steroid measurement on the same animals. For immunohistochemistry, brain hemispheres were postfixed with PFA 4% at 4°C overnight, then transferred in a 30% sucrose solution for 3 days before mounting in OCT compound and stored frozen at -20°C until slicing. Coronal frozen sections of a thickness of 35 μm were sliced with a cryostat (cryostat, Leica CM3050S) to obtain hippocampal sections conserved in a cryoprotectant solution (30% glycerol, 30% ethylene glycol and 40% PBS 1M) at -20°C until immunofluorescence staining. The other half of the brain was immediately submerged into an ethanol 70 pre-chilled to –60/–70 °C for snap-freezing, to further assess steroid levels. They were then stored at -80°C until sample preparation.

### Immunohistochemistry

#### GD 18.5 brains

The slices were washed twice in PBS1x under agitation, for 15 minutes each at RT. Antigen retrieval solution was prepared with DAKO1x (Agilent Technologies, S1699) and poured into the slides holders (80mL). The holders were placed inside the steamer at 80°C for 20 minutes before the slices were transferred into the DAKO solution for 30 minutes, covered with a lid to prevent evaporation. The slide holders with the slices were then removed and kept at RT for 45 minutes, followed by two washes of 15 minutes each. For the demasking steps, the slices were first subjected to 5 minutes of pure methanol (Sigma Aldrich, 179337), 30uL per slice. Then, hydrogen peroxyde solution was prepared fresh containing 20% methanol, 20% hydrogen peroxide (Sigma Aldrich, H1009) and 60% PBST1% and 30uL was pipetted directly to each slice twice, for 10 minutes each at RT. The slices were placed back into the slide holder containing PBS 1x and rinsed twice for 15 minutes each at RT under agitation. The slices were then permeabilized in PBST1% for 30 minutes at RT. Blocking buffer solution was prepared using 5% normal goat serum (Gibco, 11530526) in PBST1% and 30uL drops were pipetted on each slice. The slices were incubated in the blocking buffer at RT for 3 hours. The slices were then incubated with 30uL per slice for 48 hours at 4°C in PBST1% containing 2% normal goat serum with the following primary antibodies: rabbit anti-Ki-67 (1:1000, Lucerna Chem, ab15580), rat anti-Tbr2 (1:500, ebioscience, 14-4875-82), mouse anti-HopX (1:250, Santa Cruz, sc-398703) and rabbit anti-NeuroD1 (1:500, Proteintech, 12081-1-AP). After 48 hours of incubation, the antibodies were rinsed in PBS 1x twice for 15 minutes each and then with PBST1% for 30 minutes at RT under agitation. The slices were then incubated for 3 hours at RT with either of the following secondary antibodies in PBST1% and 2% normal goat serum: goat anti-rabbit Alexa Fluor 555 (1:1000, Life Technologies, A21428), goat anti-rat Alexa Fluor 488 (1:500, Life Technologies, A11006) and goat anti-mouse Alexa Fluor 488 (1:500, Invitrogen, A11029), 30uL per slice. Slices were then washed 10 minutes in PBS 1x at RT covered from light and incubated at RT for 30 minutes in 4,6 diamidino-2-phenylindole (DAPI, 1:1000) to stain nuclei before being rinsed again twice in PB 0.1M for 15 minutes under agitation. The slides were dried completely and mounted using mounting medium (Sigma Aldrich, 345789) and sealed carefully using coverslips. Once dry, the slides were stored at 4°C until microscopy imaging.

#### 10-week-old adult brains

One out of 6 slices containing hippocampal tissue were chosen to cover the whole dentate gyrus and used for immunostaining. After washes, an antigen retrieval solution was prepared with DAKO1x (Agilent Technologies, S1699) and added into the wells. Slice were incubated with antigen retrieval solution for 30 minutes at 37°C, followed by two washes of 15 minutes each. For the demasking steps, the slices were first subjected to 5 minutes of pure methanol (Sigma Aldrich, 179337), 30uL per slice. Then, slices were treated twice with a solution containing 20% methanol, 20% hydrogen peroxide (Sigma Aldrich, H1009) and 60% PBST1% for 15 minutes each and washed for 10 minutes each at RT. The slices were then rinsed twice with PBS 1x for 15 minutes each at RT under agitation. The slices were then permeabilized in PBST1% for 30 minutes at RT. Blocking buffer solution was prepared using 5% normal goat serum (Gibco, 11530526) in PBST1% and incubated in the blocking buffer at RT for 1 hour. Slices were then incubated 48 h at 4◦C in PBST1% containing 5% of normal goat or donkey serum with the following primary antibodies: rabbit anti-DCX (1:500, Cell Signaling Technology, 4604S), rat anti-Ki-67 (1:500, Lucerna Chem, ab15580), rabbit anti-GFAP (1:500, Dako Z0334), mouse anti-Sox2 (1:500, Millipore AB5603), and mouse anti-Hopx (1:500, Santa Cruz, sc-398703). After 48 hours of incubation, the sections were rinsed 3x 10min in PBST1% and incubated for 3h in either of the following secondary antibodies in PBST1% and 5% of normal goat or donkey serum: goat anti-rabbit Alexa-594 (1:500, Life Technologies, A11037), goat anti-rabbit Alexa-488 (1:500, Invitrogen, A11034), donkey anti-rabbit (1:500, Invitrogen A31573), donkey anti-mouse Alexa-488 (1:250, Invitrogen. A21202) and goat anti-rat Alexa Fluor 488 (1:500, Life Technologies, A11006). Sections were then rinsed 10min in PBS1x and incubated in 4,6 diamidino-2-phenylindole (DAPI, 1:1000) to reveal nuclei before being rinsed again 3x 10min in PB 0.1M. The slides were dried completely and mounted using mounting medium (Sigma Aldrich, 345789) and sealed carefully using coverslips. Once dry, the slides were stored at 4°C until microscopy imaging. SOX2/GFAP and DCX/Ki-67 stainings were performed on a separate set of sections and were therefore analyzed independently.

### Imaging and anatomical quantification

All images of the immunostained sections of hippocampus were acquired with a Nikon NI-E Spinning disk microscope with a 4X or a 20X objective. Nikon NIS Elements imaging software was used to do 3D supervised automatic image analysis. In embryos and adults’ slices, given the anatomical differences in the brain between the anterior and posterior axis, slices were categorized according to their anatomy as anterior, posterior or medial, taking the prenatal mouse brain atlas as the reference^73^ and Allen brain atlas for the adults. For each animal, at least three anterior and three posterior slices were taken containing the following regions of interest in the brain-the hypothalamus, the amygdala, the hippocampus, the DG and cortical thickness. DAPI staining was used for identification of the structures. Nikon NIS Element imaging software was used to perform 3D measurement analysis for the volumes of each of the brain regions. The measurements were averaged from the three anterior and posterior brain sections each. For slices classified as anterior, the hippocampi ROIs included the CA1, CA2 and CA3 and for DG most of the hilus was included. The fimbriae were not included in the ROIs for the hippocampus. The amygdala was delimited by taking the end of the external capsule and the piriform cortex bifurcation as reference, and including as much of the basolateral, central and basomedial nuclei as possible. For the hypothalamus, the paraventricular, suprachiasmatic and ventromedial nuclei were included with the zona incerta and the cerebral peduncle taken as the limits. For the slices classified as posterior, the same strategy was used for the hippocampus and the DG, whereas the amygdala was delimited by including mostly the medial nucleus and was smaller in size and closer to the cortex compared to the anterior amygdala. A similar approach was applied for the hypothalamus, using the supra-optic nucleus as landmark for the lateral delimitations. The number of labelled cells was counted for each section, in the entire thickness of the section. In the dentate gyrus cells expressing Ki-67, HOPX, Tbr2 or NeuroD1 were counted in an area containing the developing DG of GD 18.5 embryos. Whereas in 10-week-old mice, cell expressing Ki-67 were counted in the subgranular zone and the granule cell layer (GCL) of the DG, the cell expressing SOX2 and GFAP were counted only in the subgranular zone and the fractional volume occupied by DCX-labeled cells was measured in an area including the subgranular zone, the granule cell layer and the initial part of the molecular layer (ML). The fractional volume is the proportion of DCX^+^ voxels over the total number of voxels, analyzed throughout the entire thickness of each section and multiplied by the number of sections analyzed. The DCX labelling intensity was automatically detected by Nikon NIS Elements imaging software in the entire thickness of the slice within the DG. Dendritic maturation of DCX-positive cells was assessed by quantifying dendritic intersections at two defined levels of the dentate gyrus: within the granule cell layer and at the border between the granule cell layer and the molecular layer. All anatomical and cellular analyses were performed using Nikon NIS-Elements software by an experimenter blinded to the experimental groups.

### Statistical analyses

Statistical analyses for behavioral, anatomical, cellular, and hormone quantification experiments were performed using GraphPad Prism 9 (GraphPad Software, San Diego, CA, USA). Data are presented as mean ± SEM. Statistical significance was set at p < 0.05. For comparisons among IUP groups (0M, 1M, and 2M), data were analyzed separately by sex using one-way ANOVA when assumptions of homogeneity of variance were met, followed by Tukey’s multiple comparisons test. When equality of variance was not met, Brown-Forsythe ANOVA followed by Dunnett’s T3 multiple comparisons test was used. For comparisons between two groups, unpaired two-tailed t-tests were used. For prenatal hormone manipulation experiments, vehicle, DHT, and Flutamide groups were compared separately by sex. Treatment effects were analyzed using one-way ANOVA or Brown-Forsythe ANOVA, as appropriate, followed by post-hoc comparisons of DHT and Flutamide groups against the vehicle group. For single-nucleus RNA-sequencing analyses, statistical methods are described in the corresponding sections. Differential gene expression was assessed using MAST with Benjamini-Hochberg correction for multiple testing. Cell-subtype proportion differences were assessed using quasibinomial generalized linear models followed by Benjamini-Hochberg correction. Trajectory-dependent gene expression was analyzed using tradeSeq, with adjusted p-values calculated using the Benjamini-Hochberg method.

---

**Supplementary Figure 1.**
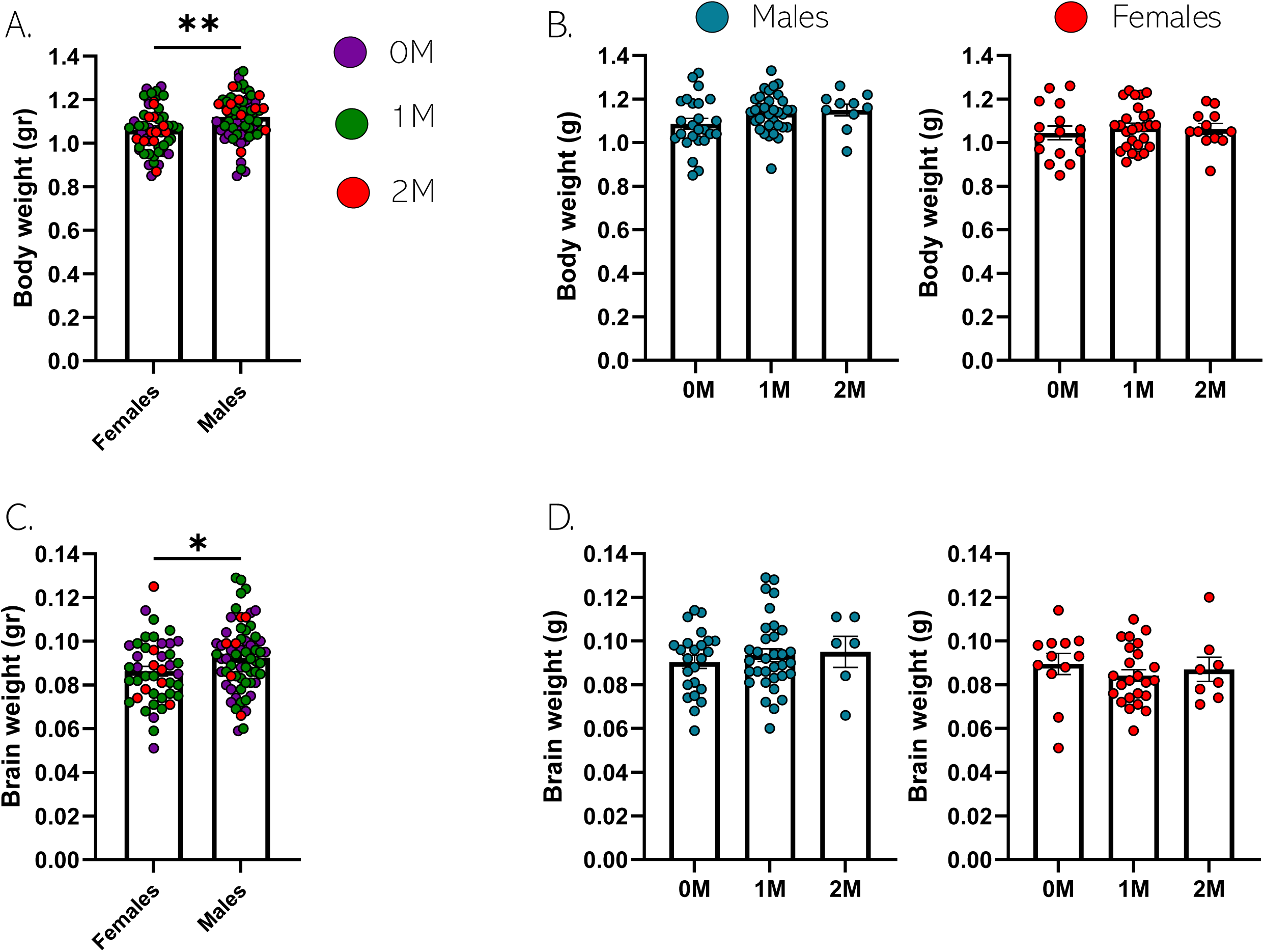
Effect of IUP and sex on body and brain weight at GD 18.5: **A.** Histogram of body weight in females and males (t121=3.124, p = 0.02, unpaired t test, two tailed, n = 55 and 68 per group). **B.** Histogram of body weight in males (left) and females (right) according to IUP (**Males**: F2,65 = 2.228, p = 0.116, one-way ANOVA, n = 25, 33 and 10 mice per group. **Females**: F2,52 = 0.305, p = 0.738, one-way ANOVA, n = 16, 27 and 12 per group). **C.** Brain weight in females and males (t105=2.056, p = 0.042, unpaired t test, two tailed, n = 44 and 63 per group). **D.** Brain weight in males (left) and females (right) according to IUP (**Males**: F2,60 = 0.356, p = 0.701, one-way ANOVA, n = 24, 34 and 6 mice per group. **Females**: F2,41 = 0.542, p = 0.586, one-way ANOVA, n = 12, 24 and 8 per group). Histograms show average ± SEM, * p<0.05, ** p<0.01, ***p<0.001, ns = not significant.

**Supplementary Figure 2.**
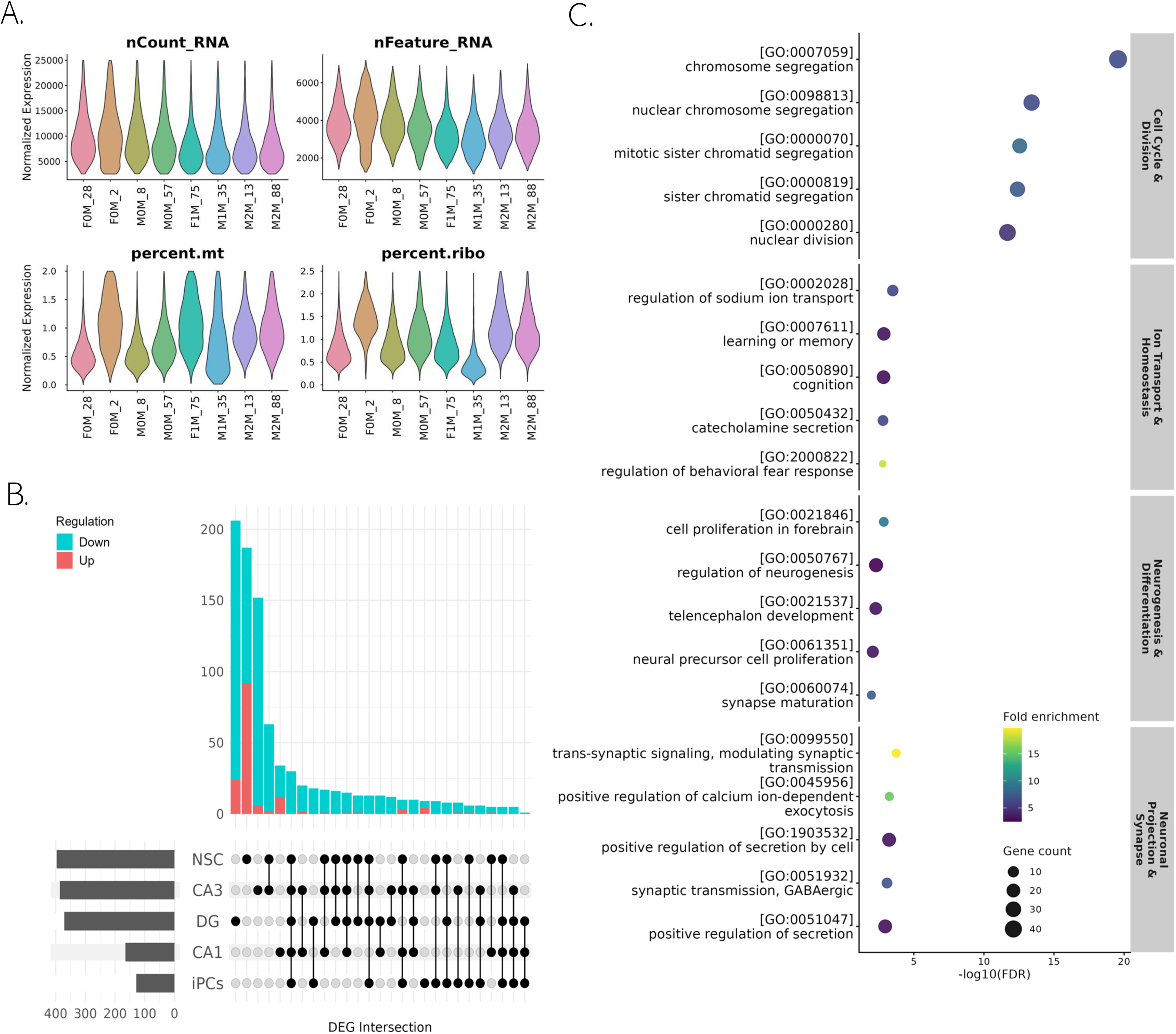
**A.** QC metrics for nuclei included in snRNAseq analysis. **B.** Upset plot of differentially expressed genes identified for each cell type comparison. **C.** Top five GO terms from each major category identified from differentially expressed genes among 1M/2M vs. 0M in NSCs.

**Supplementary Figure 3.**
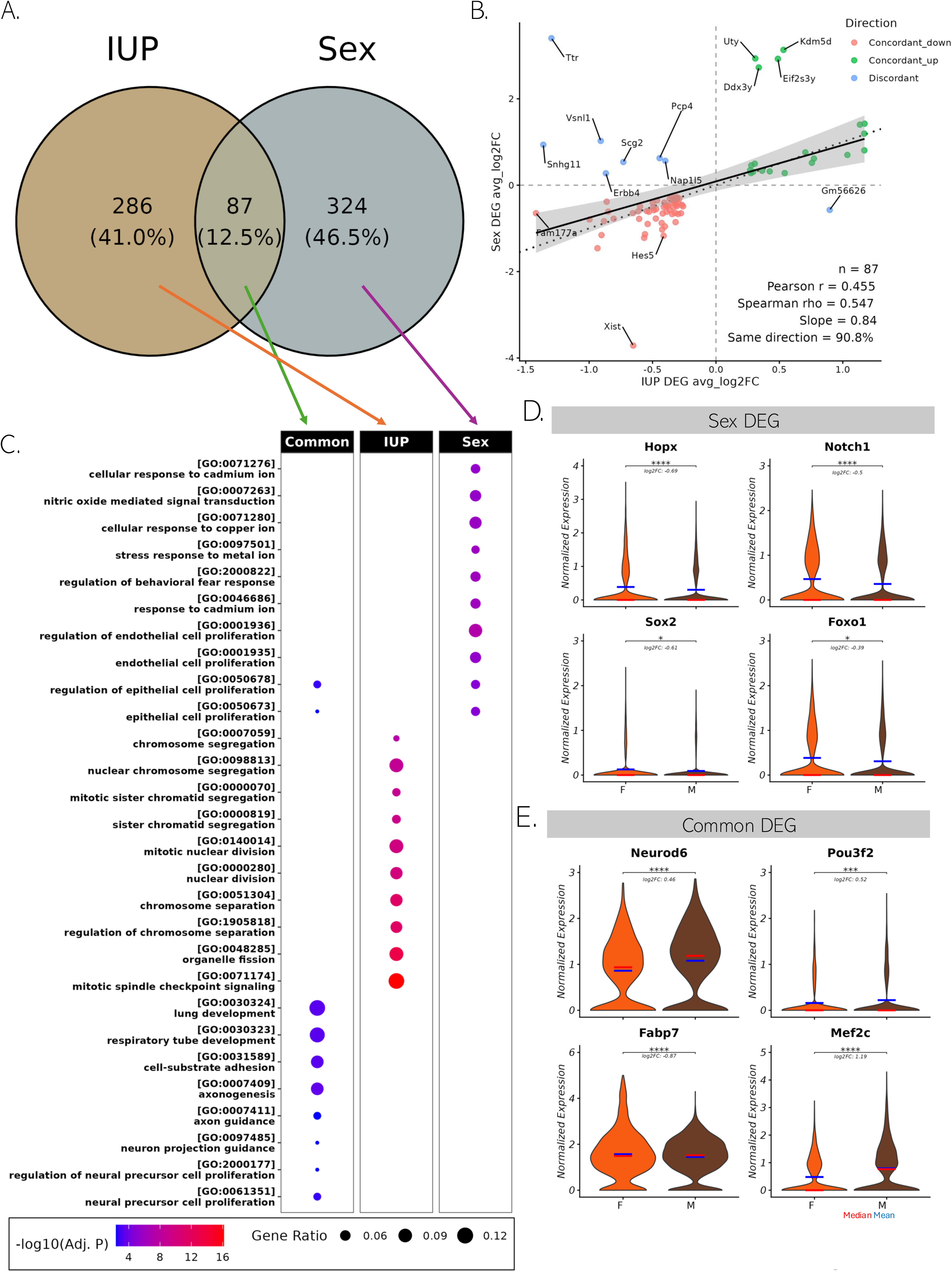
**A.** Venn diagram comparing differentially expressed genes identified in comparisons of IUP (1M/2M vs. 0M) and sex (M vs. F). **B.** Comparison of fold changes of differentially expressed genes in common between the two comparisons. **C.** Full set of GO terms identified from differentially expressed gene expression analyses. **D.** Violin plots showing normalized expression of representative sex-specific DEGs in NSCs, including Hopx, Notch1, Sox2 and Foxo1. **E.** Violin plots showing normalized expression of representative DEGs common to the IUP and sex comparisons in NSCs, including Neurod6, Pou3f2, Fabp7 and Mef2c.

**Supplementary Figure 4.**
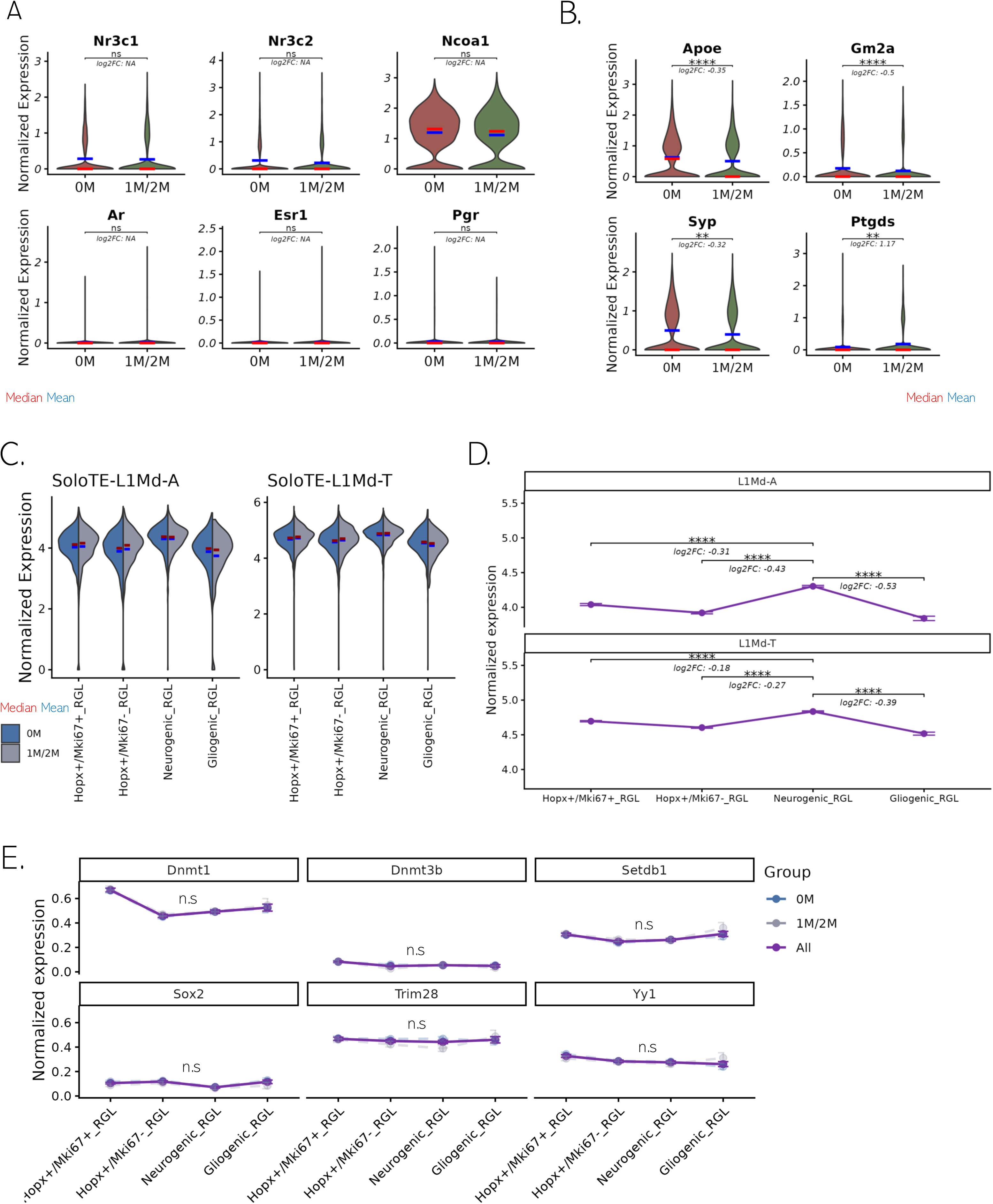
**A.** Violin plots showing normalized expression of steroid hormone receptor genes and **(B)** additional neural stem cell related genes. **C-D.** Expression of LINE-1 subfamilies of interest from SoloTE, comparing among NSC subtypes (MAST, Padj < 0.05). **E.** Expression of known LINE-1 regulator genes across NSC subtypes (MAST, Padj < 0.05).

**Supplementary Figure 5.**
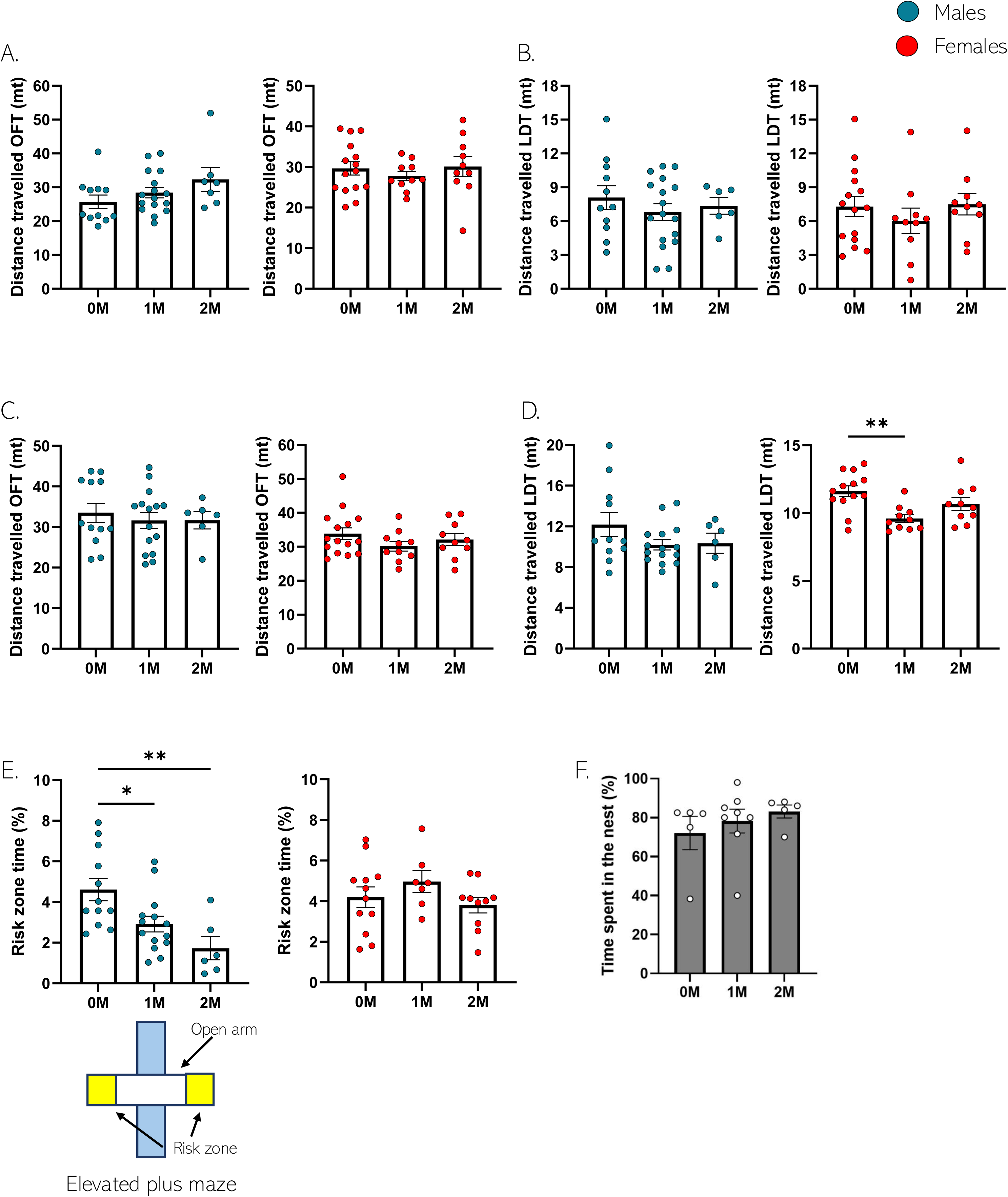
Effect of IUP on locomotor activity and anxiety. Maternal behavior of CD1 mothers: **A.** Distance travelled in an OFT at PND22 by males (left panel) and females (right panel) (**Males**: F2,31 = 1.892, p = 0.168, one-way ANOVA, n = 11, 16 and 7 mice per group. **Females**: F2,32 = 0.435, p = 0.650, one-way ANOVA, n = 15, 10 and 10 per group). **B.** Distance travelled in an LDT at PND23 by males (left panel) and females (right panel) (**Males**: F2,31 = 0.583, p = 0.564, one-way ANOVA, n = 11, 17 and 6 mice per group. **Females**: F2,32 = 0.576, p = 0.567, one-way ANOVA, n = 15, 10 and 10 per group). **C.** Distance travelled in an OFT at PND55 by males (left panel) and females (right panel) (**Males**: F2,30 = 0.238, p = 0.789, one-way ANOVA, n = 12, 15 and 6 mice per group. **Females**: F2,32 = 1.249, p = 0.300, one-way ANOVA, n = 15, 10 and 10 per group). **D.** Distance travelled in an LDT at PND56 by males (left panel) and females (right panel) (**Males**: F2,29 = 1.645, p = 0.210, one-way ANOVA, n = 11, 15 and 6 mice per group. **Females**: F2,30 = 6.671, p = 0.004, one-way ANOVA, n = 13, 10 and 10 per group; Tukey’s multiple comparisons test, 0M vs 1M: p = 0.003, 0M vs 2M: p = 0.212; 1M vs 2M: p = 0.184). **E.** Percentage of time spent in the risk zone in an EPM by males (left panel) and females (right panel) (**Males**: F2,29 = 7.109, p = 0.003, one-way ANOVA test, n = 12, 14 and 6 mice per group; Tukey’s multiple comparisons test, 0M vs 1M: p = 0.033, 0M vs 2M: p = 0.004; 1M vs 2M: p = 0.304. **Females**: F2,26 = 1.424, p = 0.259, one-way ANOVA, n = 12, 7 and 10 per group). **F.** Time spent in the nest by CD1 foster mothers (F2,15 = 0.611, p = 0.556, one-way ANOVA, n = 5, 8 and 5 per group). Histograms show average ± SEM, * p<0.05, ** p<0.01, ns = not significant.

**Supplementary Figure 6.**
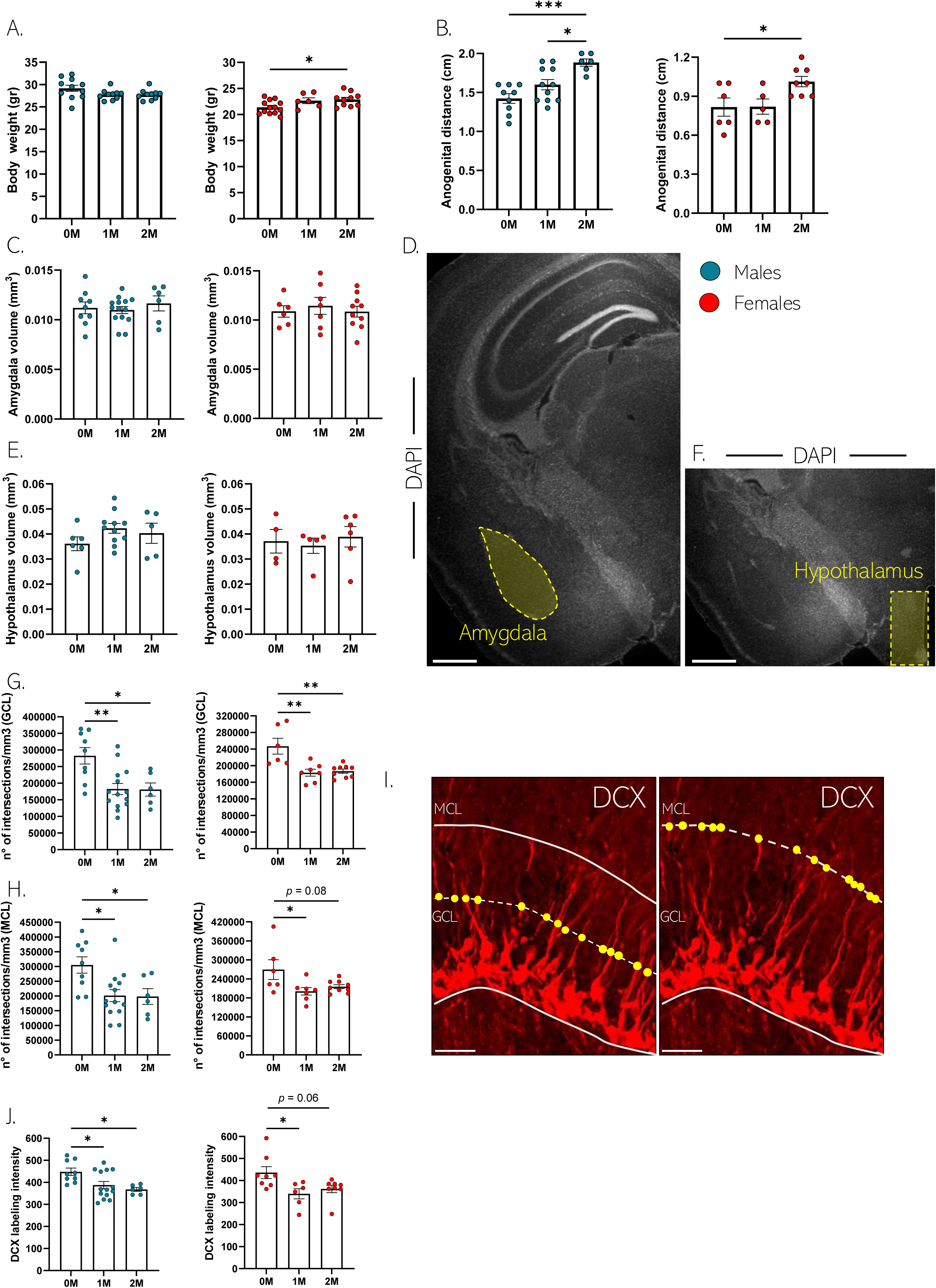
Effect of IUP on anatomy and immature neurons maturation: **A.** Body weight in males (left panel) and females (right panel) at 10-week-old (**Males**: F2,28 = 2.261, p = 0.123, one-way ANOVA test, n = 11, 10 and 10 mice per group. **Females**: F2,26 = 4.155, p = 0.027, one-way ANOVA test, n = 13, 6 and 10 per group; Tukey’s multiple comparisons test, 0M vs 1M: p = 0.147, 0M vs 2M: p = 0.031; 1M vs 2M: p = 0.938). **B.** Anogenital distance in males (left panel) and females (right panel) at 10 weeks of age (**Males**: F2,23 = 10.69, p = <0.001, one-way ANOVA test, n = 9, 11 and 6 mice per group; Tukey’s multiple comparisons test, 0M vs 1M: p = 0.179, 0M vs 2M: p = 0.179; 1M vs 2M: p = >0.999. **Females**: F2,26 = 4.155, p = 0.027, one-way ANOVA test, n = 13, 6 and 10 per group; Tukey’s multiple comparisons test, 0M vs 1M: p = 0.114, 0M vs 2M: p = <0.001; 1M vs 2M: p = 0.019). **C.** Analysis of amygdala volume in males (left panel) and females (right panel) (**Males**: F2,26 = 0.370, p = 0.694, one-way ANOVA test, n = 9, 14 and 6 mice per group. **Females**: F2,20 = 0.239, p = 0.789, one-way ANOVA test, n = 6, 7 and 10 per group). **D.** Representative confocal microscopy image of the amygdala (highlighted in yellow) immunostained for DAPI (scale bar 1000µm). **E.** Analysis of hypothalamus volumes in males (left panel) and females (right panel) (**Males**: F2,19 = 1.455, p = 0.258, one-way ANOVA test, n = 6, 11 and 5 mice per group. **Females**: F2,12 = 0.217, p = 0.808, one-way ANOVA test, n = 5, 5 and 6 per group). **F.** Representative confocal microscopy image of the hypothalamus (highlighted in yellow) immunostained for DAPI (scale bar 1000µm). **G.** Histograms showing the number of DCX^+^ dendrites intersections in the GCL of the DG in males (left panel) and females (right panel) (**Males**: F2,26 = 7.618, p = 0.002, one-way ANOVA test, n = 9, 15 and 6 mice per group; Tukey’s multiple comparisons test, 0M vs 1M: p = 0.003, 0M vs 2M: p = 0.016; 1M vs 2M: p = 0.999. **Females**: F2,19 = 10.00, p = 0.001, one-way ANOVA test, n = 6, 7 and 9 per group; Tukey’s multiple comparisons test, 0M vs 1M: p = 0.002, 0M vs 2M: p = 0.002; 1M vs 2M: p = 0.967). **H.** Histograms showing the number of DCX^+^ dendrites intersections in the MCL of the DG in males (left panel) and females (right panel) (**Males**: F2,26 = 5.819, p = 0.008, one-way ANOVA test, n = 9, 14 and 6 mice per group; Tukey’s multiple comparisons test, 0M vs 1M: p = 0.003, 0M vs 2M: p = 0.016; 1M vs 2M: p = 0.999. **Females**: F2,19 = 10.00, p = 0.001, one-way ANOVA test, n = 6, 7 and 9 per group; Tukey’s multiple comparisons test, 0M vs 1M: p = 0.010, 0M vs 2M: p = 0.035; 1M vs 2M: p = 0.997). **I.** Confocal maximal projection of DG slices immunostained for DCX (in red) highlighting the number of DCX^+^ intersections (yellow dots) in the GCL (left panel) and MCL (right panel) (scale bar 10µm). **J.** DCX labeling intensity in DG slices of males (left panel) and females (right panel) (**Males**: F2,26 = 5.227, p = 0.012, one-way ANOVA test, n = 9, 14 and 6 mice per group; Tukey’s multiple comparisons test, 0M vs 1M: p = 0.032, 0M vs 2M: p = 0.020; 1M vs 2M: p = 0.724. **Females**: F2,19 = 4.855, p = 0.020, one-way ANOVA test, n = 8, 6 and 8 per group; Tukey’s multiple comparisons test, 0M vs 1M: p = 0.025, 0M vs 2M: p = 0.066; 1M vs 2M: p = 0.794). Histograms show average ± SEM, * p<0.05, ** p<0.01, ns = not significant.

**Supplementary Figure 7.**
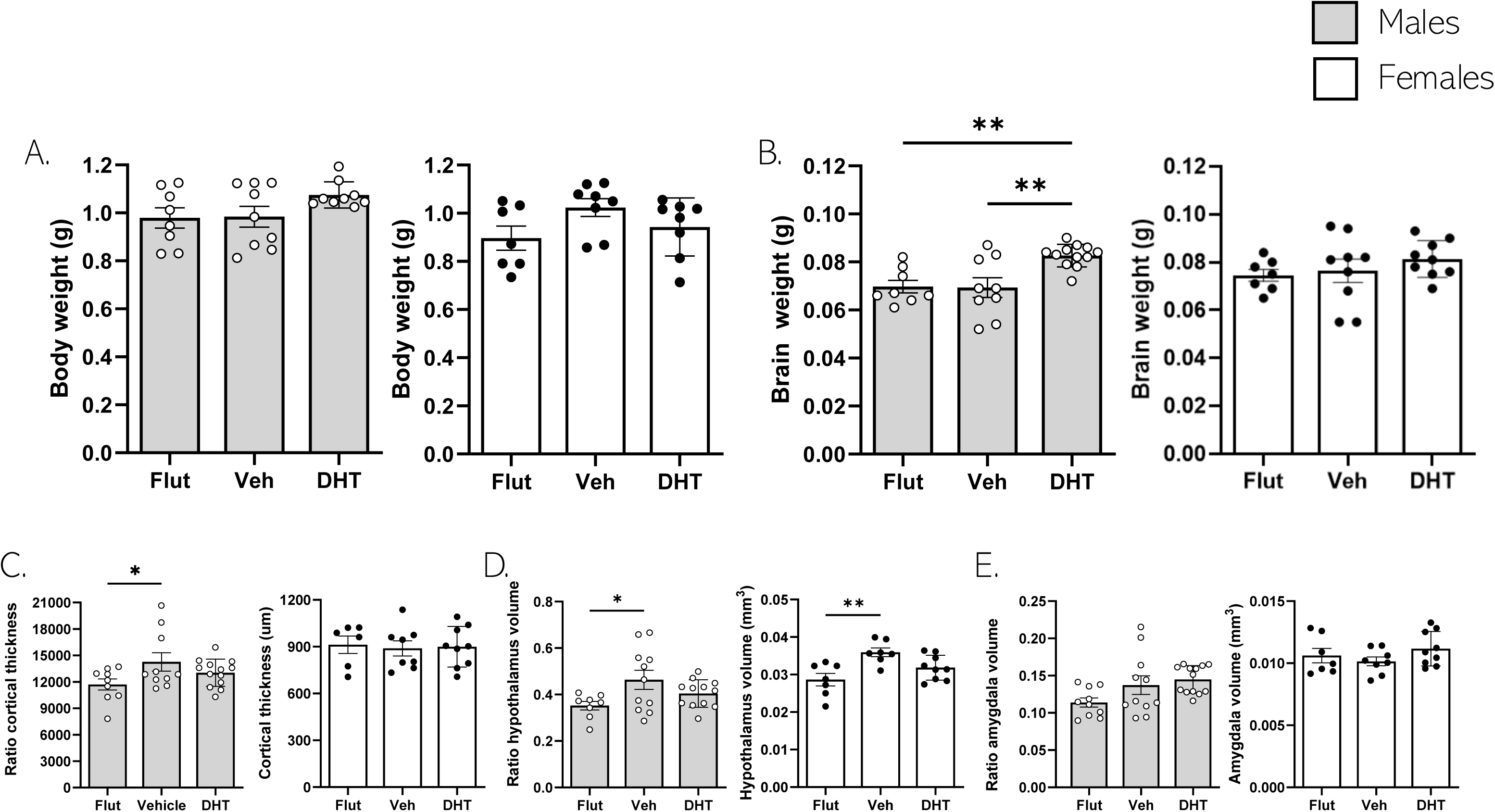
Effect of steroids on embryos’ development: **A.** Body weight in males (left panel) and females (right panel) in embryos at GD 18.5 (**Males**: F2,17.61 = 2.205, p = 0.1398, Brown-Forsythe ANOVA test, n = 8, 9 and 9 mice per group, Dunnett’s T3 multiple comparisons test with the mean of Veh group: Veh vs Flut: p = 0.9997, Veh vs DHT: p = 0.1538. **Females**: F2,20 = 2.223, p = 0.1343, one-way ANOVA test, n = 7, 8 and 8 mice per group, Dunnett’s T3 multiple comparisons test with the mean of Veh group: Veh vs Flut: p = 0.0929, Veh vs DHT: p = 0.3153). **B.** Brain weight in males (left panel) and females (right panel) in embryos at GD 18.5 (**Males**: F2,26 = 8.573, p = 0.0014, one-way ANOVA test, n = 8, 9 and 12 mice per group, Dunnett’s T3 multiple comparisons test with the mean of Veh group: Veh vs Flut: p = 0.9923, Veh vs DHT: p = 0.0026. **Females**: F2,22 = 0.8974, p = 0.4220, one-way ANOVA test, n = 7, 9 and 9 mice per group, Dunnett’s T3 multiple comparisons test with the mean of Veh group: Veh vs Flut: p = 0.9162, Veh vs DHT: p = 0.5304). **C.** Histograms of cortical thickness in males (left panel) and females (right panel) at GD 18.5 (**Males**: F2,29 = 2.915, p = 0.0402, one-way ANOVA test, n = 9, 10 and 14 mice per group, Dunnett’s T3 multiple comparisons test with the mean of Veh group: Veh vs Flut: p = 0.9997, Veh vs DHT: p = 0.1538. **Females**: F2,20 = 2.223, p = 0.1343, one-way ANOVA test, n = 7, 8 and 8 mice per group, Dunnett’s T3 multiple comparisons test with the mean of Veh group: Veh vs Flut: p = 0.0408, Veh vs DHT: p = 0.3532). **D.** Histograms of hypothalamus volumes in males (left panel) and females (right panel) at GD 18.5 (**Males**: F2,17.37 = 4.450, p = 0.0275, Brown-Forsythe ANOVA test, n = 8, 11 and 12 mice per group, Dunnett’s T3 multiple comparisons test with the mean of Veh group: Veh vs Flut: p = 0.0319, Veh vs DHT: p = 0.2552. **Females**: F2,20 = 7.228, p = 0.0043, one-way ANOVA test, n = 7, 7 and 9 mice per group, Dunnett’s T3 multiple comparisons test with the mean of Veh group: Veh vs Flut: p = 0.0022, Veh vs DHT: p = 0.0633). **E.** Histograms of amygdala volumes in males (left panel) and females (right panel) at GD 18.5 (**Males**: F2,18.21 = 3.435, p = 0.0542, Brown-Forsythe ANOVA test, n = 10, 11 and 14 mice per group. **Females**: F2,21 = 1.270, p = 0.3016, one-way ANOVA test, n = 7, 8 and 9 mice per group). Histograms show average ± SEM, * p<0.032, **p<0.0021, ns = not significant.

